# Mapping the Neural Dynamics of Locomotion across the *Drosophila* Brain

**DOI:** 10.1101/2022.03.20.485047

**Authors:** Luke E. Brezovec, Andrew B. Berger, Shaul Druckmann, Thomas R. Clandinin

## Abstract

Walking is a fundamental mode of locomotion, yet its neural correlates are unknown at brain-wide scale in any animal. We use volumetric two-photon imaging to map neural activity associated with walking across the entire brain of *Drosophila*. We detect locomotor signals in approximately 40% of the brain, identify a global signal associated with the transition from rest to walking, and define clustered neural signals selectively associated with changes in forward or angular velocity. These networks span functionally diverse brain regions, and include regions that have not been previously linked to locomotion. We also identify time-varying trajectories of neural activity that anticipate future movements, and that represent sequential engagement of clusters of neurons with different behavioral selectivity. These motor maps suggest a dynamical systems framework for constructing walking maneuvers reminiscent of models of forelimb reaching in primates and set a foundation for understanding how local circuits interact across large-scale networks.

## Introduction

Topographic maps have provided fundamental insights into the functional organization of the brain. Such maps of neural activity have revealed spatial order in sensory systems, motor control, and cognitive processing in many different animals (Kaas 1997; Leyton and Sherrington 1917; Patel et al. 2014). The existence of these functional maps reflects the fact that neural processing is often distributed across large populations of cells whose spatial arrangements reveal coding principles of the system. For example, functional maps of motor cortex in primates have revealed how movements of different limbs are spatially segregated, a necessary step to revealing how specific movement trajectories are encoded by the dynamics of neural populations (Churchland et al., 2012; Graziano and Aflalo, 2007; Penfield and Boldrey, 1937; Shenoy et al., 2013). Many other brain regions contain signatures of behavioral movements such as walking (Ferreira-Pinto et al., 2018; Grillner and El Manira, 2020), and yet the brain-wide functional topography of walking behavior has not been described in any animal.

Walking subserves a diverse array of behavioral goals in many animals, and must be shaped by both sensory inputs and internal states. Despite its central role, locomotor control is incompletely understood (Merel et al., 2019; Schwartz et al., 2016; Straka et al., 2018). In vertebrates, many brain regions play key roles in shaping locomotion, including the basal ganglia, brainstem, cerebellum, motor cortex, and spinal cord (Ferreira-Pinto et al., 2018; Grillner and El Manira, 2020; Kiehn and Dougherty, 2013). More recently, broad swaths of both sensory and non-sensory cortex have been shown to contain motor-related signals (Clancy et al., 2019; Kaplan and Zimmer, 2020; Karadimas et al., 2020; Musall et al., 2019; Stringer et al., 2019; Zatka-Haas et al., 2021). In addition, brain-wide signatures of neural activity associated with swimming and crawling have been measured in larval zebrafish, larval *Drosophila*, and C. elegans (Ahrens et al., 2012; Chen et al., 2018; Dunn et al., 2016; Hallinen et al., 2021; Kato et al., 2015; Kim et al., 2017; Mu et al., 2020; Naumann et al., 2016; Nguyena et al., 2015; Susoy et al., 2021). Moreover, recent work has described whole-brain imaging techniques in adult flies in the context of state dependent changes in activity and metabolism (Aimon et al., 2019; Mann et al., 2017; Mann et al., 2021; Schaffer et al., 2021; Tainton-Heap et al., 2021), as well as sensory evoked responses (Harris et al., 2015; Münch et al., 2021; Pacheco et al., 2021). Given the accumulating evidence across species for widespread distribution of motor signatures, directly measuring this topography is likely to provide insight into how the brain produces walking behavior.

Insects have long provided valuable insights into the biomechanics and neural control of locomotion (Cruse et al., 2009; Hughes, 1952; Manton, 1973; Wilson, 1966). More recently, the scalability of *Drosophila* behavioral measurements has allowed a variety of quantitative descriptions of the structure of walking dynamics in flies (Berman et al., 2014; Branson et al., 2009; Chun et al., 2021; DeAngelis et al., 2019; Kain et al., 2013; Katsov et al., 2017; Mendes et al., 2013; Strauss and Heisenberg, 1990). Furthermore, the majority of descending neurons (DNs) that relay movement commands from the central brain to the ventral nerve cord (*Drosophila*’s equivalent of a spinal cord) have been identified (Hsu and Bhandawat, 2016; Namiki et al., 2018). Functional measurements and perturbations of these populations have identified cell types that can specifically induce forward and backward walking (Bidaye et al., 2014; Bidaye et al., 2020; Cande et al., 2018; Rayshubskiy et al., 2020). Moreover, characterization of circuits that relay visual information to DNs has provided insights into how vision can monitor self-motion and guide steering control (Suver et al., 2016). In addition, targeted studies of specific visual interneurons have revealed how efference-copy signals are relayed into the visual system to modulate processing (Chiappe et al., 2010; Cruz et al., 2021; Fujiwara et al., 2017; Kim et al., 2015; Kim et al., 2017; Maimon et al., 2010; Suver et al., 2012). Finally, navigation related neural signals in the central complex, as well as associative learning signals in the mushroom body, are also modulated by walking (Cohn et al., 2015; Fisher et al., 2019; Weir and Dickinson, 2015; Zolin et al., 2021). However, how this cornucopia of motor signals might be spatially organized across and within brain regions, and coordinated in time as walking occurs, is unknown.

Here, we develop a volumetric two-photon imaging and analysis pipeline to extract neural activity from across the entire *Drosophila* brain as the animal behaves. We then describe a volumetric registration technique that allows signals to be quantitatively compared across brain regions and individuals. Using this method, we explore the brain-wide neural dynamics of walking. We discover that locomotor signals are widespread, extending throughout approximately 40% of the brain volume, and account for the dominant dimensions of neural activity. We discern both a global state change signal that defines the transition between moving and not moving, as well as particular brain regions that contain information specific to the forward or rotational velocity of the fly. Moreover, we observe activity in some brain regions that precedes changes in velocity by at least 300 milliseconds, while activity in other regions lags behind behavior by more than a second. The temporal evolution of this activity thus describes a stereotyped pattern of recruitment of specific brain regions during walking. Combining the fine spatial structure of the topographic map of neural selectivity within individual brain regions with its temporal sequence of activation suggests a dynamical systems framework for relating neural activity to specific behavioral maneuvers. Taken together, these studies identify a brain-wide spatiotemporal topography of walking, setting a critical foundation for relating signals in specific, genetically targeted circuits and cell types to global dynamics.

## Results

### A novel method for whole-brain imaging in walking *Drosophila*

We sought to map neural activity associated with walking behavior across the *Drosophila* brain, to quantitatively compare these signals across individuals, and to develop mathematical models that relate neural and behavioral measurements. To do this we developed a pipeline for measuring neural activity across the whole volume of the brain while recording the animal’s locomotion in the dark (Figure 1A). We expressed two fluorescent indicators in all neurons: GCaMP6f to monitor neural activity, and myristylated-tdTomato as a structural marker (Chen et al., 2013; Pfeiffer et al., 2010). Flies were head-fixed and the posterior head cuticle was removed to expose the brain, including most of the optic lobe neuropils and the central brain (Figure S1). We then employed two-photon imaging with resonant scanning to achieve a volume imaging rate of 1.8 Hz, collecting approximately 1.6M voxels per volume, each occupying 2.6 × 2.6 × 5 µm. Signals from both fluorophores were acquired simultaneously. During each 30 minute imaging session the animal’s walking trajectory was measured by recording the rotations of an air-suspended treadmill ball. At the end of each recording, we collected a high spatial resolution (0.6 × 0.6 × 1 um/voxel) anatomical scan of the tdTomato signal. These structural measurements allowed us to register every voxel of neural activity across individuals with high spatial accuracy. Because individual flies display non-linear anatomical differences, we iteratively applied both affine alignment as well as a diffeomorphic warping algorithm employed in human fMRI (Advanced Normalization Tools, ANTs (Avants et al., 2009; Avants et al., 2011)) to produce a “mean” brain (Figure S1). This process created a common space in which data from across all flies, and across all brain regions were well aligned (Figure 1B). To parse these data into anatomically defined brain regions, we registered a canonical atlas of labeled neuropils (Ito et al., 2014; Jenett et al., 2012) into our common space using a two-step process that combined ANTs with a neural network, SynthMorph, trained to generalize over contrast variance, as seen in this type of multimodal image registration problem (Hoffman et al., 2020).

**Figure 1.**
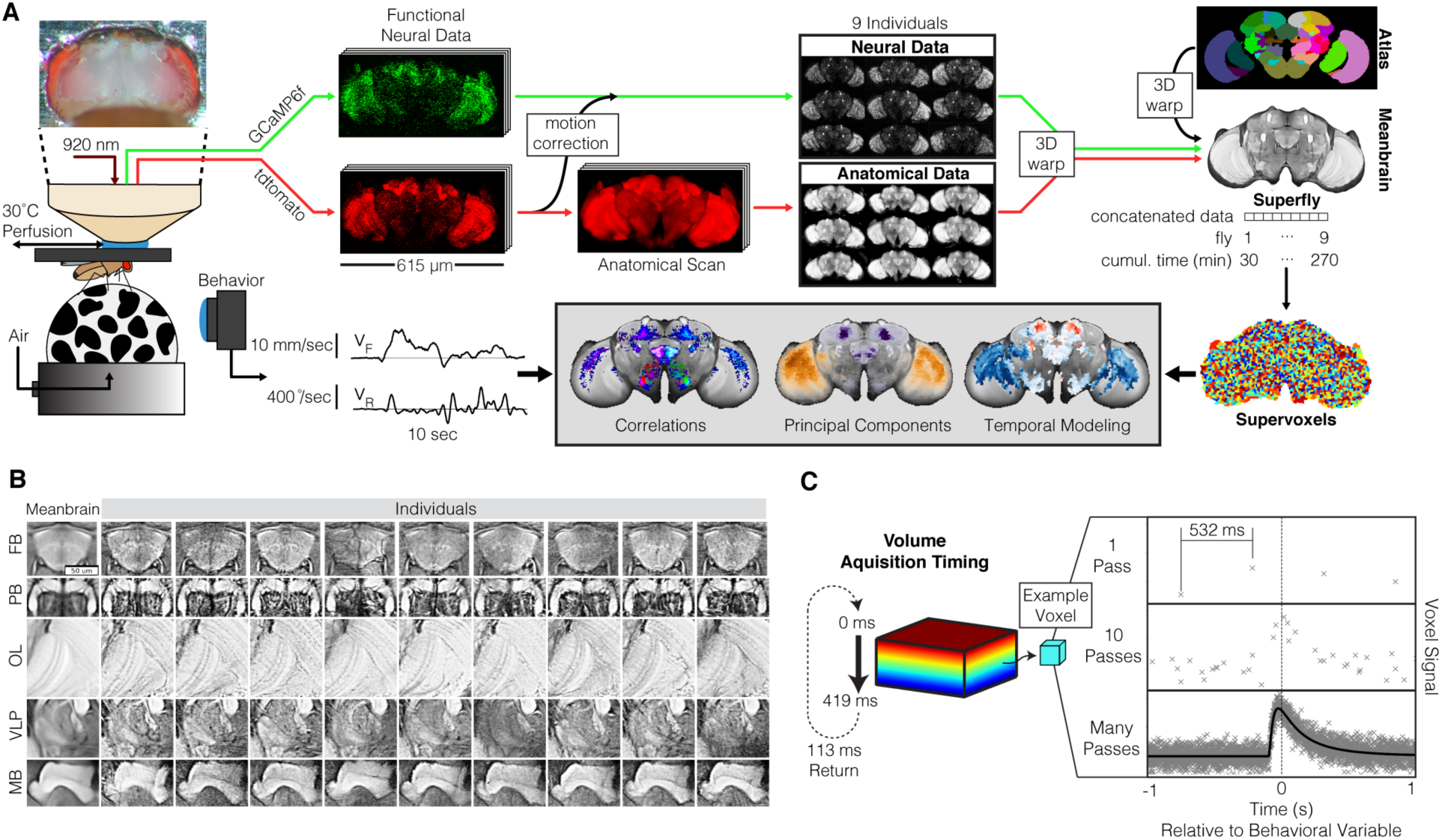
Whole-brain imaging in walking *Drosophila*. (A) Overview of pipeline. After dissection of the posterior head cuticle, the fly is mounted under a two-photon microscope and walks on an air-suspended ball in the dark. GCaMP6f is expressed pan-neuronally, as is a structural marker, tdTomato. Volumes were acquired at 1.8 Hz at a resolution of 2.6 × 2.6 × 5um for 30 minutes to capture neural activity; a subsequent anatomical scan was taken at higher spatial resolution (0.6um x 0.6um x 1um). The structural marker was used to correct brain motion. Nine individuals passed all quality control metrics (see Supplemental Figure 1) and were used for subsequent analyses. All nine datasets were registered into a single mean brain using the structural marker; these warp parameters were then applied to the functional neural data to bring all flies into the same space. A standard anatomical atlas was also registered into this common space. Concatenating all data resulted in a “Superfly” with 270 cumulative minutes of neural and behavioral recording. Voxels were clustered into supervoxels (see methods). Behavior was collected at 50 Hz and walking trajectories were decomposed into forward and rotational velocities. Data was analyzed using correlations, principal components, and temporal modeling. (B) Qualitative comparison of alignment quality. The meanbrain is compared with anatomical scans from each individual. Five representative regions of the brain are cropped to the same coordinates after alignment. (C) Schematic illustration of the logic for obtaining high temporal resolution filters from low temporal resolution data. Temporal acquisition of each voxel is well defined. A single pass through a behavior of interest yields a poor estimate of the underlying filter; with many passes at different offsets a high resolution filter can be measured.

By registering all our data into a common space and concatenating the neural and behavioral signals across flies (and sessions), we created a unified dataset (the “superfly”) where 1.6M voxels have each been sampled 30,456 times over the course of 4.5 hr and 9 individuals (Figure 1A). For each of these individuals, we sampled the movement of the ball as a proxy for walking at a temporal frequency of 50Hz. These measurements were then temporally smoothed and aligned with neural activity data from each voxel with a precision of approximately 5 ms (see Methods). The large number of temporal samples in our unified neural activity dataset, combined with these precise timestamps on the acquisition of each voxel relative to behavior, allowed us to construct temporal filters that relate neural activity to behavior across the dataset (Figure 1C). In most subsequent cases, we also reduced the number of features in the neural activity dataset by agglomerative clustering of neighboring voxels with similar responses to create “supervoxels” (Figure 1A). In the following analyses, we describe the relationship between neural activity across the brain and walking behavior using combinations of correlations, principal components, and temporal modeling.

### Neural encoding of velocity space is widespread across the brain

In darkness, tethered flies spontaneously initiate bouts of walking activity composed of sequences of turns and straight runs (spanning seconds), separated by periods of quiescence and grooming (lasting tens of seconds). We reasoned that, under these conditions, neural activity might reveal a common pattern of spatiotemporal dynamics, including signals that initiate movement, execute specific maneuvers, and relay information about ongoing movement to sensory systems. To relate neural activity to behavior, we decomposed the walking trajectories of each fly in the dataset into forward velocity (V_F_) and rotational velocity (V_R_ (right turn, clockwise), and V_L_ (left turn, counterclockwise)) (Figure 2A). Then, we calculated the correlation of each voxel’s neural activity with each of these three components (Figure 2B). To visualize the spatial structure of these correlations, we colored the correlation with each velocity component as an axis in red-green-blue (RGB) color space (Figure 2C). We observed that signals were widespread, with 39% of the brain volume correlating with at least one of the three behavioral variables (p<0.001, Bonferroni Corrected; Figure S2). V_F_ correlations exhibited strong mirror symmetry across the midline, and consisted of voxels that correlated only to V_F_ (15% brain volume) as well as voxels that correlated with multiple variables (20% brain volume). In contrast, V_R_ and V_L_ correlations were anti-symmetric, with high levels of correlation on the side of the brain that is ipsilateral to the direction of the turn, with significantly fewer correlated voxels on the contralateral side. In other words, turning to the left (counterclockwise) is strongly correlated with activity in the left hemisphere, and vice versa for a right turn. Most voxels that correlated with V_R_ or V_L_ also correlated with V_F_ (20% brain volume), with relatively few correlating only with turning (4% brain volume). In all, 90% of brain voxels that correlated with behavior displayed a tuning preference in velocity space, with only 10% of voxels responding indiscriminately to all velocity components. Taken together, these data argue for the presence of distributed neural signals that relate to specific velocity features of locomotion.

**Figure 2.**
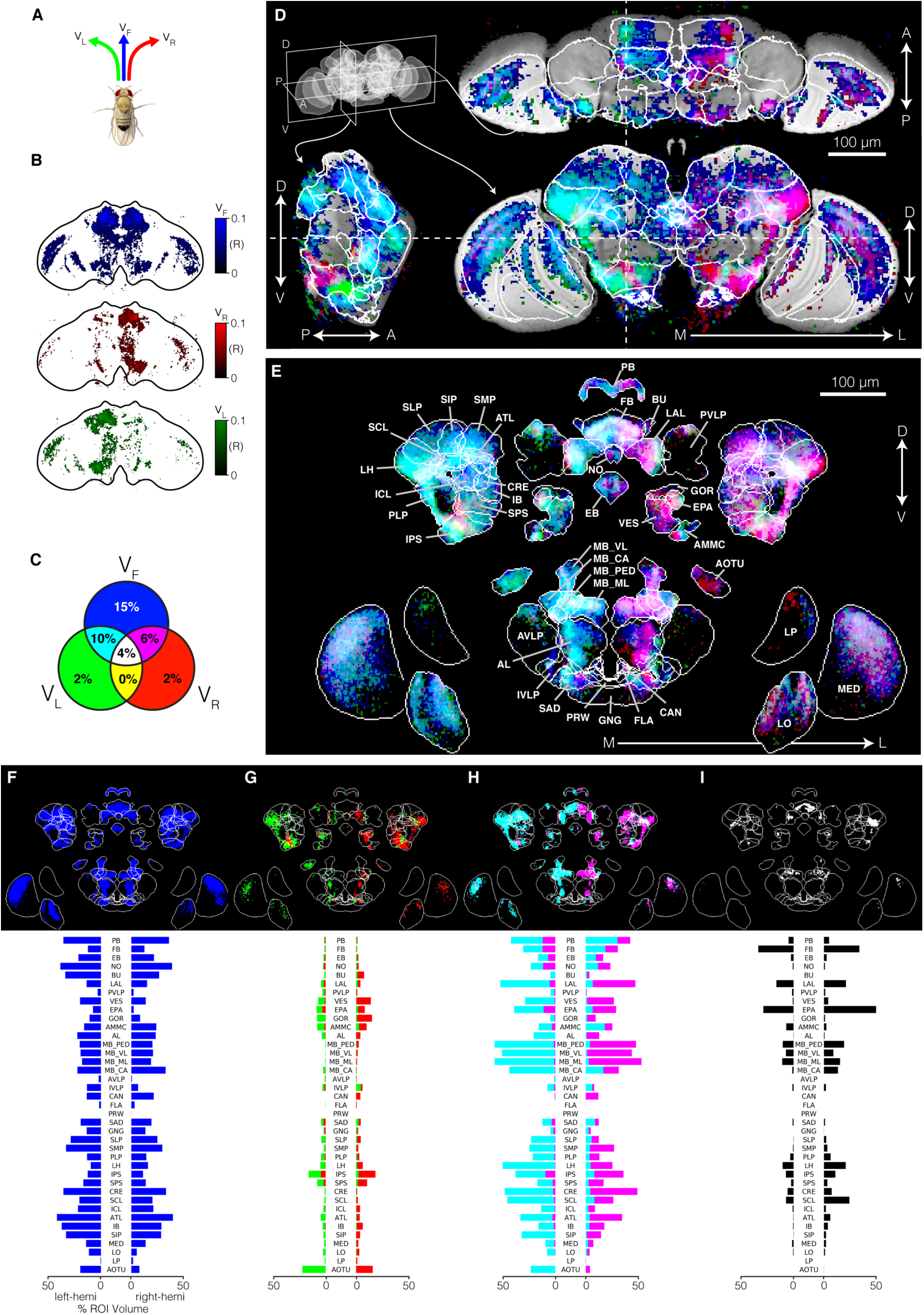
Neural encoding of movement velocity is widespread across the brain and highly structured. (A) The trajectory of the fly was decomposed into a forward velocity, as well as positive (clockwise) and negative (anticlockwise) angular velocities. (B) Correlation map of individual voxels for each behavioral velocity. A single example slice is shown. Voxels above a significance threshold were not considered for analysis and were set to white (p > 0.0001; see Methods). (C) Venn diagram distribution of voxel-tuning types. Voxels were categorically assigned to a velocity set. Numbers indicate percent of voxels in each category. Red, green and blue correspond to a neural signal that correlated with only one of the three velocity variables, while cyan, magenta, and yellow correspond to correlations with two of the variables, and white corresponds to a correlation with all variables. (D) Example slices through orthogonal planes of the brain. RGB values were independently set based on correlation with velocity components. (E) Maximum intensity projection of the partially exploded mean brain illustrating the categorical assignment of voxel tuning (following the color code in (C). (F-I) As in (E), separating categorical voxel types into separate panels. Bar charts quantify percent of voxels in each anatomical ROI that fall into each velocity category.

We next wondered how these voxel correlations map onto the anatomical substructures of the brain. Visualizing individual slices through the volume along three orthogonal axes reveals spatially structured correlation patterns, spanning much of the central brain and optic lobes (Figure 2D; Figure S2). However, a 3D representation of this data proved challenging, given the intricate anatomy of the brain. In contrast, cutting the brain into its anatomically defined regions and visualizing them separately allowed the correlations to be visualized well, but the relationships between signals in neighboring neuropils was lost. We therefore split the brain into large regions and then took a maximum intensity projection through each substructure (Figure 2E). This segmentation revealed that behaviorally correlated voxel signals were highly spatially structured, with some signals being restricted to specific anatomical regions, and others revealing additional structure in which specific layers or sub-regions of a given anatomical region display selective behavioral correlations.

Although almost all neuropils contained signals that were correlated with behavior, their velocity preferences were highly non-uniform across brain regions (Figure 2E-I). To examine the structure of these voxel categories across each anatomical region, we next calculated the fraction of the volume of each anatomical region that was composed of each voxel category. As expected, the central complex navigation and premotor region (including the Protocerebral Bridge (PB), Fan-shaped Body (FB), Ellipsoid Body (EB) and Nodulus (NO)) was well represented. Similarly, areas that are associated with extensive innervation by descending neurons (DNs), and hence are likely involved in motor control, were also highly engaged, with strong signals in the Inferior Posterior Slope (IPS), Superior Posterior Slope (SPS), Vest (VES), Superior Medial Protocerebrum (SMP), and Superior Lateral Protocerebrum (SLP). Notably, however, other regions that also include significant descending neuron innervation are not strongly correlated with walking, suggesting that they may be predominantly engaged in controlling other motor behaviors. Finally, we also observe significant behavioral signals in higher order sensory areas, including signals related to visual processing (the Anterior Optic Tubercle (AOTU), the Lobula (LO), and the Medulla), olfactory processing (Antenna Lobe (AL) and Lateral Horn (LH)), and auditory information (Antennal Mechanosensory and Motor Center (AMMC)). Finally, the associative learning center, the mushroom body (MB), is also strongly correlated with walking behavior. Taken together, these data demonstrate that locomotor behavior is associated with changes in neural activity across navigation areas, motor areas, higher order sensory areas and association areas, revealing the extensive impact of movement on neural processing.

### Distinct topographic maps of locomotor signals exist in multiple brain regions

This initial characterization revealed that signals associated with specific combinations of forward and angular locomotor velocities were differentially mapped across many brain areas (Figure 2D-I). For example, the Inferior Posterior Slope (IPS), the Anterior Optic Tubercle (AOTU) and the Vest (VES) have a significant fraction of their volumes occupied by the relatively rare signals associated only with angular velocity (“turning only”); the Mushroom Body (MB) and Lateral Accessory Lobe (LAL) are dominated by the forward-and-turning type; and, the Superior Medial Procerebrum (SMP) and the Superior Intermediate Protocerebrum (SIP) had mostly the forward-only type. We also observed that although ipsilateral hemispheres were well-correlated with turning, there were also notable exceptions, most obviously within subregions of the Inferior Posterior Slope (IPS) and Lateral Accessory Lobe (LAL), in which contralateral signals were correlated with their respective turn direction.

Given this rich distribution of voxel tuning properties, we next examined whether different voxel signals were spatially organized within each neuropil. Taking slices through orthogonal planes of each neuropil, we observed clear spatial organization at the sub-neuropil scale (Figure 3; Figure S3). The Posterior Lateral Protocerebrum (PLP) contained an approximately 15 micron diameter column of voxels running along the dorsal-ventral axis which uniformly correlated with both forward and ipsilateral turning velocities (Figure 3A). The Inferior Posterior Slope (IPS) contained discrete compartments correlating with either left or right turns, and were anti-symmetric between hemispheres (Figure 3B). The Lobula (LO) displayed segregated bands of correlation, consistent with known anatomy, that had mixed selectivity for forward velocity and ipsilateral turning (Figure 3C). The Lateral Horn (LH) contained a dorsal-ventral split, with the dorsal half correlating with forward velocity, and the ventral half correlating with both forward velocity and ipsilateral turning (Figure 3D). The Fan-shaped Body (FB) contained two distinct layers with identical velocity correlations (forward velocity and ipsilateral turning) (Figure 3E). The entire Protocerebral Bridge (PB) correlated with forward velocity, in addition to an overlaid alternating pattern of left and right turning correlations, matching known functional connectivity (Figure 3F) (Franconville et al., 2018; Green et al., 2017). The Nodulus (NO) was uniformly correlated with forward velocity, but contained distinct glomeruli that correlate with contralateral turning (Figure 3G). Similar to the IPS and the LH, the Lateral Accessory Lobe (LAL) contained distinct compartments, all correlated with forward velocity, but with the medial portion of the neuropil correlated with ipsilateral turns, and a lateral portion correlated with contralateral turns (Figure 3H). In summary, our experimental and analytical approach has revealed topographic maps containing extensive functional order at the sub-neuropil level, relating neural activity to specific changes in behavior.

**Figure 3.**
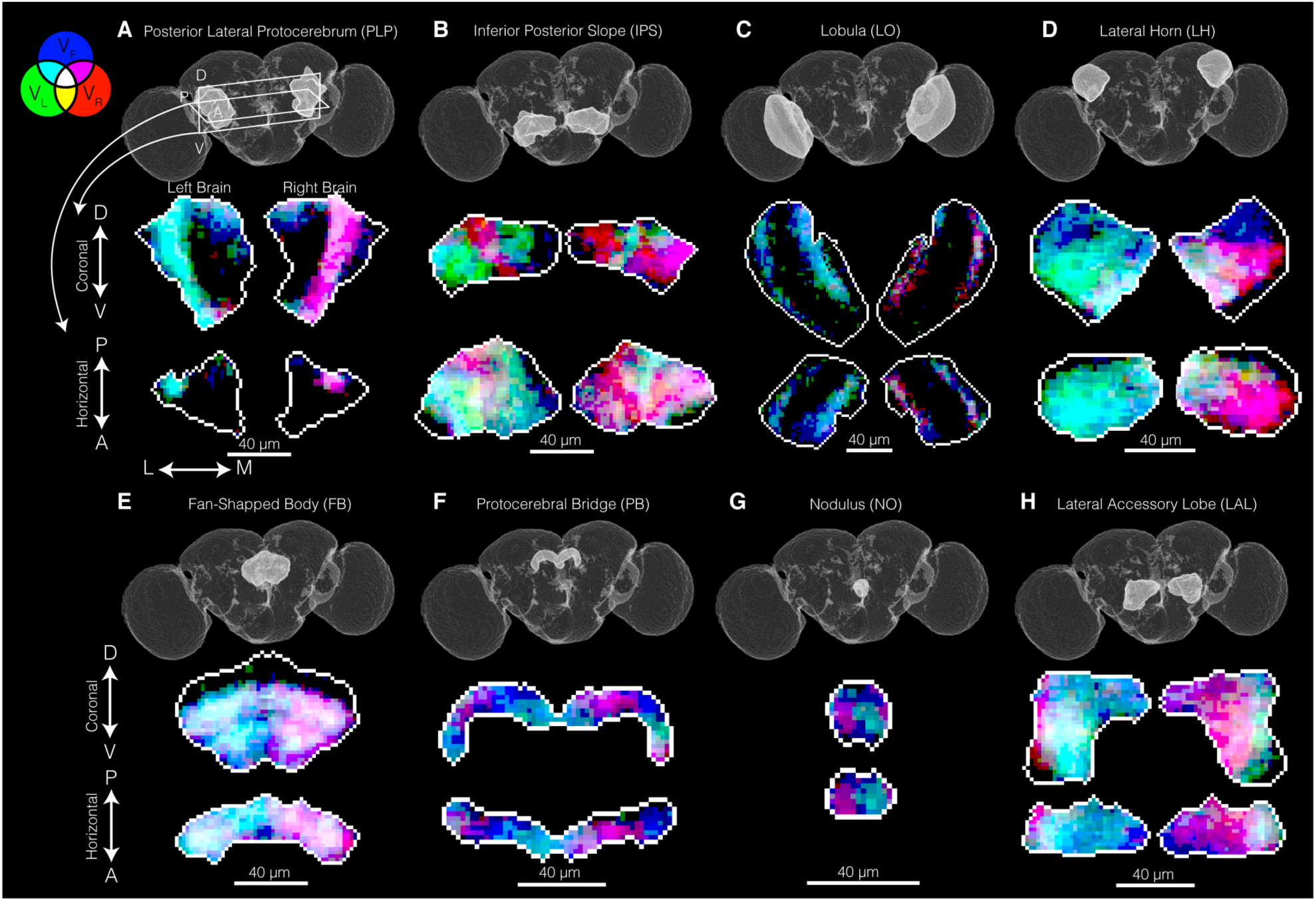
Topographic maps of neural tuning to behavioral velocities in specific brain regions. Coronal and horizontal slices shown. (A) Posterior Lateral Protocerebrum (PLP). (B) Inferior Posterior Slope (IPS). (C) Lobula (LO). (D) Lateral Horn (LH). (E) Fan-shaped Body (FB). (F) Protocerebral Bridge (PB). (G). Ellipsoid Body. (H). Lateral Accessory Lobe (LAL). Color coding as in Figure 2. Related to Supplemental Figure 3.

### Neural activity in the central brain is highly predictive of locomotor velocity

Our analysis of correlations between voxel activity and behavior considered each voxel independently. We next sought to consider all voxels simultaneously in order to identify shared, common signals, and to use these signals to predict the structure of behavior. To accomplish this, we first reduced the number of features (corresponding to the number of voxels) by agglomerative clustering in which individual voxels were sequentially merged into “supervoxels” so as to minimize the variance of each cluster. This clustering step also served to increase the signal to noise ratio (Figure S4). As an initial glimpse into the temporal structure of these data, we then performed Principal Component Analysis (PCA) on the full neuronal (but not behavioral) dataset (30,456 timepoints, 98,000 spatial supervoxels).

This analysis revealed that the most significant dimensions of variation were highly structured (Figure 4A). To explore the relationship between the first three principal components and behavior, we projected neural activity onto each of the eigenvectors to calculate their temporal signals, and then binned the signals in 2-D and 1-D velocity spaces (Figure 4B,C). Strikingly, these dimensions were highly structured by the relationship with behavior, as described by changes in forward and angular velocities. In particular, the value of PC1 was low when the fly was not moving and high otherwise, while PC2 increased with forward velocity but was invariant with respect to angular velocity, and PC3 encoded information about the sign of the angular velocity. To quantify this observation, we set a velocity threshold to approximate the transition from being stopped to being walking, and then calculated the correlation between each principal component and each velocity component, both below and above the threshold (Figure 4C,D). As expected, the under-threshold correlations were high for PC1 and low for PC2 and PC3, while PC2 displayed the highest correlation for above-threshold forward velocity, and PC3 showed opposite correlations to above-threshold left and right angular velocities. Consistent with these average correlations, similar relationships were observed when comparing the values of each PC overlaid with the corresponding behavioral variables in a single bout (Figure 4E). Finally, in addition to these first three PCs having distinct correlations with behavior, we observed that many other PCs were highly structured in velocity space (Figure S4). Thus, these data demonstrate that there are signatures of locomotor behavior that are widely shared across the brain.

**Figure 4.**
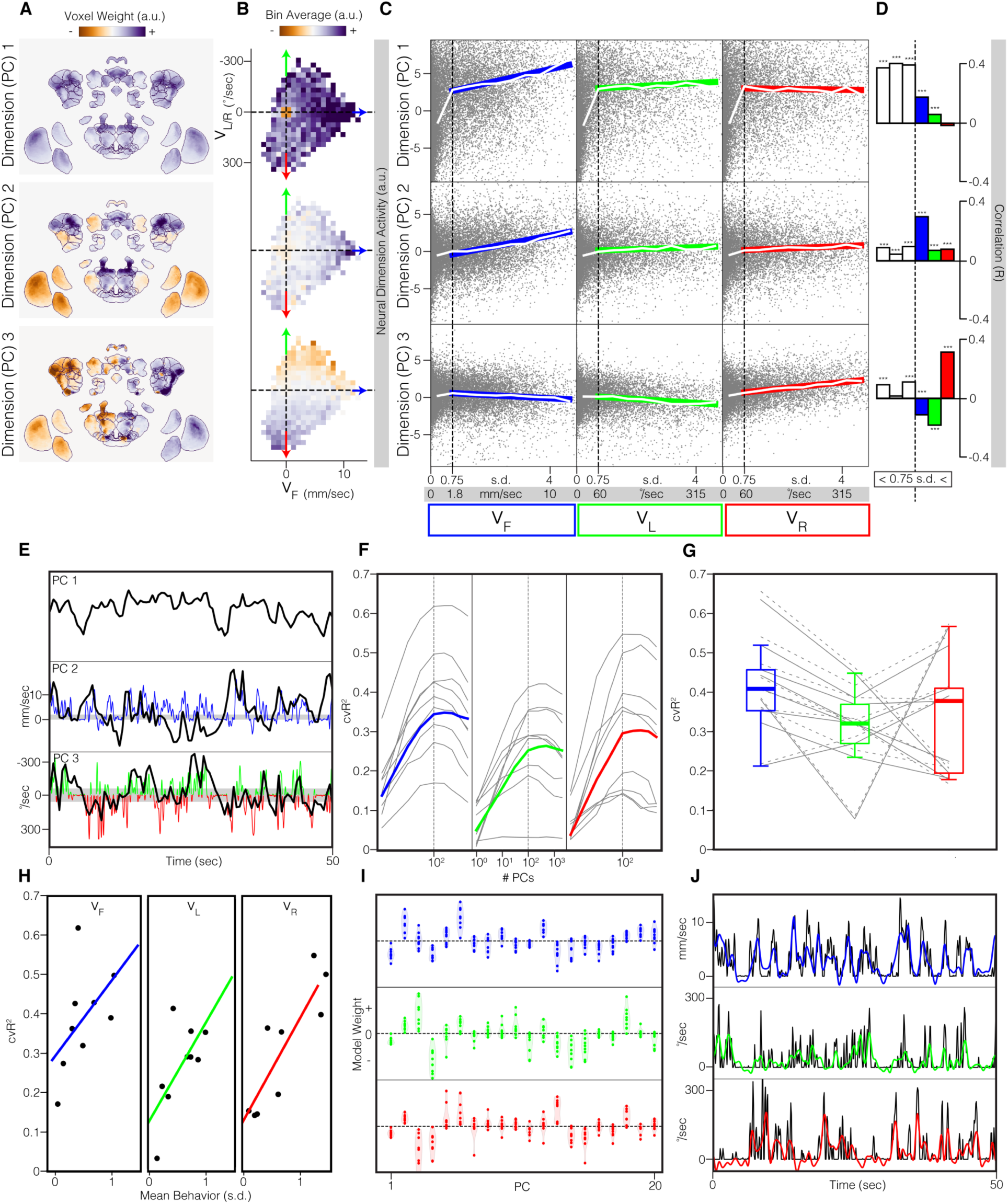
Low dimensions of neural activity are structured and predictive of velocity space. (A) The first three principal components of the integrated whole-brain neural dataset (n = 9 flies). Maximum intensity projections are colored by voxel weights for each principal component. (B) Neural data projected onto each principal component and binned in 2D velocity space. We note that the small skew in the angular velocity distribution reflects a small rotational asymmetry in placement of each fly on the ball. (C) 1D velocity spaces versus principal component values. Grey dots are single timepoints of a collected neural volume. Vertical dashed black line represents the chosen threshold between moving and not-moving. Colored lines denote the linear regression of data above this threshold. White lines are mean bin values, with a width of 0.5 s.d.. Vertical white lines represent ± 1 SEM, but are too small to be visible. (D) Quantification of data in (C). Correlations were measured independently for below and above the movement threshold. *p < 0.05, **p < 0.01, ***p 0.001. (E) Example trace of first 3 PCs and behavior. Black lines are PCs, blue is forward velocity, and red and green are rotational velocities. PCs are overlayed with relevant behaviors. Gray shading shows 0.75 s.d. threshold used in previous panel. (F) Cross-validated R^2^ prediction accuracy on test data for linear models independent fit to predict velocity components. For each velocity component, separate models were fit using different numbers of principal components as input features (x-axis). Curving gray lines are individual flies. Colored lines are the integrated superfly. Vertical dashed gray lines represent the number of components used in subsequent panels. (G) Cross-validated R^2^ prediction accuracy for a model with 100 input principal components. Solid gray lines are individual flies predicted using the principal components from the integrated superfly dataset, while the dashed gray lines are individual flies fit using their respective individual principal components. Box plots represent first quartile, median, and third quartile, while whiskers are 1.5 times the interquartile-range. (H) Same data as solid gray lines in (F), but compared with the mean amount of each velocity component seen in each individual fly. Colored lines are linear regression. (I) Model weights fit for the first 20 principal components. Each dot is an individual fly. (J) Actual (black) versus predicted (color) velocity traces. Principal components and behavior were interpolated to 10 Hz for this visualization.

To test the relationship between these dimensions of neural activity and behavior, we built a linear model that used PCs as input variables and predicted forward and angular velocities as outputs. We first determined how many PCs were necessary to maximize prediction accuracy, and observed that model performance improved over the first 100 PCs used, before declining as additional PCs were incorporated (Figure 4F). Notably, all three behavioral variables displayed comparable predictability on held-out test data, with average R^2^ values of approximately 0.3, a highly significant prediction capability for a model that generalized across flies (Figure 4G). Moreover, comparing model predictions on individual animals revealed that flies that moved less along a given velocity component were poorly predicted (R^2^ of approximately 0), while those that explored a velocity component more were well predicted (with R^2^ values up to 0.6) (Figure 4H). Significantly, despite differences in prediction accuracy, the weights assigned to each PC for each fly, and for each behavior, were highly stereotyped such that each individual fly weighted the same dimension of neural activity comparably (Figure 4I). Additionally, the weights assigned to the first three PCs were in line with our previous interpretations: PC1 was used to predict all velocity components, PC2 was predominantly used to predict forward velocity, and PC3 was used for left and right turning predictions, flipping sign between the two. Finally, at a qualitative level, the predicted velocity traces from these models closely tracked those seen in the behavioral measurements for both forward and angular velocities (Figure 4J). Taken together, these analyses demonstrate that we have identified a common neural activity space that predicts locomotor behavior across flies.

### Demixing the unique contribution of each velocity component to neural activity

The structure of walking behavior produces correlated changes in forward and angular velocities over a wide range of timescales (Berman et al., 2014; Katsov et al., 2017). Such behavioral correlations can obfuscate the relationship between neural activity and a specific behavioral variable. To estimate the effect of such correlations, we built a linear model that predicted neural activity in each voxel using four separate behavioral variables, namely a binary variable describing locomotor state (walking or stopped), forward velocity, and left and right angular velocities (see Methods). Using this approach we distinguished between the variance of each voxel that could be explained by several behavior variables, versus the *unique* variance of the voxel that could be explained only by one specific variable. That is, we identified neural activity signals that could not be accounted for using any of the other three behavior variables, and was therefore unique, on a voxel by voxel basis.

Using this approach, we reconstructed brainwide maps of the contributions of each voxel to each single variable trained separately, as well as of the unique contributions of each of the four behavioral variables, trained together (Figure 5A). This comparison revealed that the brain-wide maps of unique contributions identified sparser subsets of voxels than the single variable models (Figure 5A). For example, while the single variable forward walking and locomotor state (stopped versus walking) maps were very similar, the maps created by demixing revealed that forward velocity was encoded by a spatially organized subset of voxels, while locomotor state was not encoded uniquely anywhere in the brain.

**Figure 5.**
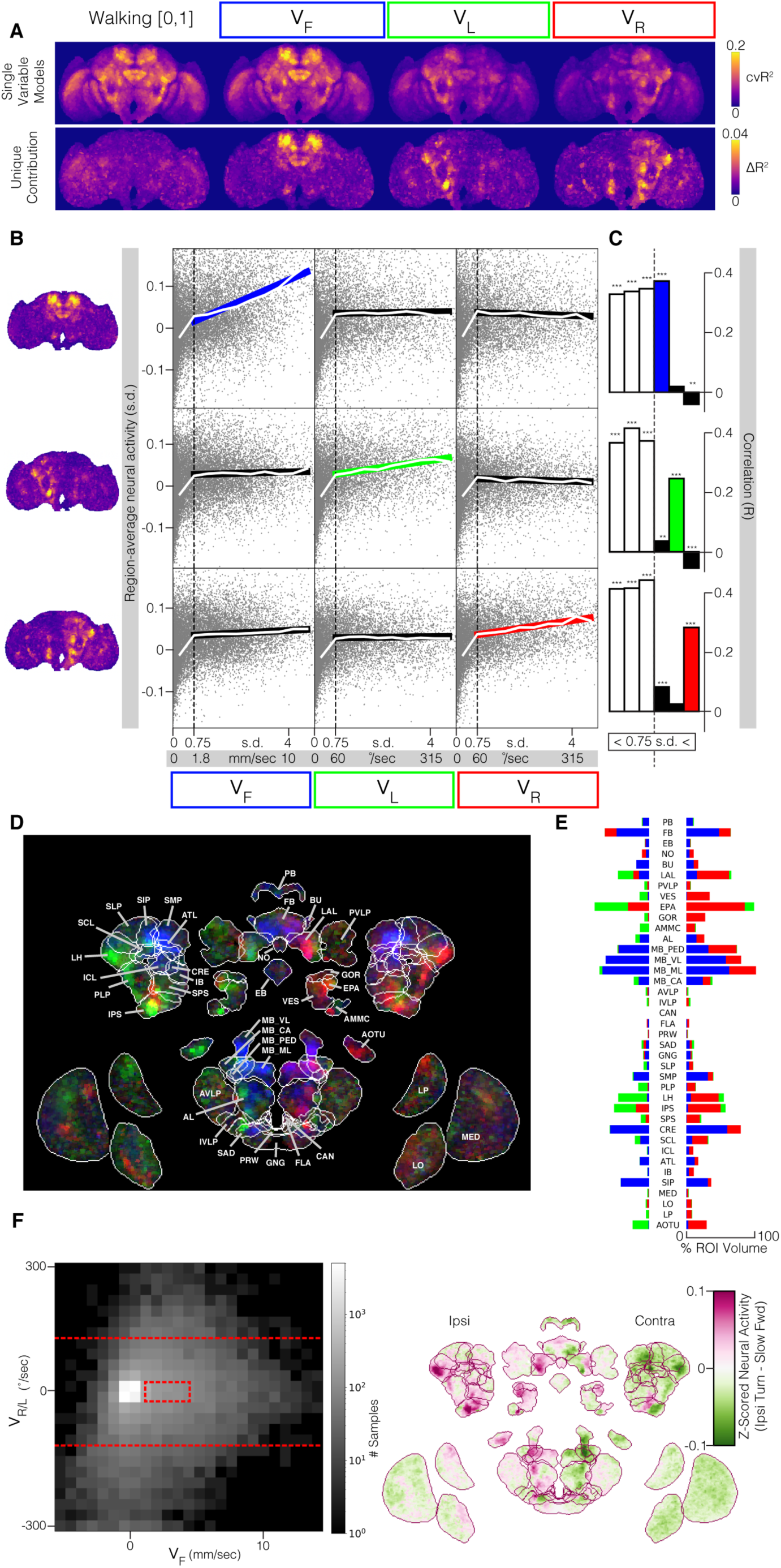
Demixing the unique contributions of each velocity component to neural activity using linear models. (A) Cross-validated linear models fit to predict voxel activities given various behavior input features. Maximum projections of R^2^ prediction accuracy. Top row: models are given only one behavior variable. Bottom row: the unique contribution of each behavior variable is measured by comparing a model that receives all four behavior variables to one that receives all variables except for the variable of interest. (B) Relationship between mean neural activity across identified functional regions from (A) versus each behavioral variable. Grey dots are single timepoints. Vertical dashed black line represents the threshold between moving and not-moving. Colored lines and thick black lines are the linear regression of data above threshold. White lines are mean bin values, with width of 0.5 s.d.. Vertical white lines represent ± 1 SEM, but are too small to be visible. (C) Quantification of data in (B). Correlations were measured independently below and above the movement threshold. *p < 0.05, **p < 0.01, ***p 0.001. (D) Maximum intensity projection colored by functional regions defined in (A). (E) Percent of voxels in each anatomical ROI that fall into each functional region. (F) Difference in brain-wide activity between a fly walking forward slowly versus turning. Red box indicates bounds for slow walking; red lines indicate lower thresholds for turning. Maximum intensity projection; average of (left turn - slow) and (right turn - slow) after mirroring the brain across the midline.

As the unique contributions of forward velocity and left and right angular velocities revealed clearly distinct spatial networks, we wanted to more closely examine these functional regions and their tuning properties. To do this, we averaged neural activity across each of the functional brain regions, and plotted their relationships to each of the three behavioral variables (Figure 5B). As before, we set a velocity threshold to approximate the speed at which a fly transitions between walking and not walking. Then, we calculated the correlation between each functional region and each velocity component, separately for below and above the threshold (Figure 5C). Significantly, we found that all three regions were highly correlated when the fly was stopped (below the threshold), strongly arguing for a wide-spread increase in neural activity associated with the initiation of movement (Figure S5). Moreover, when moving (above the threshold), each brain region had a different response to each of the three velocity variables, and as expected, each region had strong correlations with only one behavioral velocity component. Overall, this analysis has allowed us to spatially define functional brain regions that relate to unique axes of velocity space.

By visualizing these functional regions, we observed that both the correlation maps (Figure 2), and the PC maps (Figure 4), were very similar to the demixed maps (Figure 5D,E, Figure S5). In particular, the demixed forward velocity functional region closely matches both PC2, and the forward correlation map. Similarly, the demixed left and right angular velocity functional regions closely match PC3, as well as the left and right turning correlation maps. Finally, this demixing analysis revealed an additional feature of the relationship between neural activity and angular velocity that could not be discerned from either our correlation maps, or our PCA-based approach. In particular, the demixing linear model revealed that when the fly is turning, neural activity on the ipsilateral hemisphere increases in specific regions, while the contralateral hemisphere experiences a corresponding decrease in activity in the same regions (Figure 5F). Thus, initiating a turn engages widespread, reciprocal changes in neural activity across ipsi- and contra-lateral hemispheres. Overall, our analyses demonstrate that three distinct approaches, namely correlations (Figure 2), principal components (Figure 4), and linear models (Figure 5), all converge in their description of the relationship between brain-wide neural activity and locomotion.

### The temporal relationship between neural activity and behavior varies across brain regions

We next sought to determine the temporal relationship between neural activity and behavior. To do this, we cross-correlated neural activity of each supervoxel with each of the three behavioral variables (Vf, Vr and Vl), sweeping a range of relative time offsets between the two signals. At each time offset, we calculated the correlation between each supervoxel and each behavioral variable (Figure 6A) and plotted the time of peak correlation for each supervoxel (Figure 6B-D). The results revealed strong spatiotemporal structure across the brain, identifying brain regions whose signals were correlated with future changes in behavior, regions whose maximum correlations were contemporaneous with behavior, and regions that were most correlated with changes in behavior that had happened in the past. At a high level, the earliest correlations between neural activity and behavior emerged in specific neuropils approximately 300 milliseconds before the change in behavior, with the latest peak correlations persisting for more than 1 second after the change in behavior. Moreover, these spatiotemporal patterns were dramatically different for forward and rotational velocities, consistent with our instantaneous analyses, again suggesting distinct patterns of brain regions are engaged with forward and rotational velocities. In addition, we observed a positive correlation between the time of peak correlation and the width of the temporal kernel, i.e. supervoxels that contain delayed information about behavior displayed a broad temporal kernel, while supervoxels that contain anticipatory information about behavior displayed narrow temporal kernels (Figure 6E-G). Thus, lagging signals do not appear to capture as much temporal precision as leading signals, and instead reflect integration over a longer history of behavior. Finally, we emphasize that these time intervals reflect peak correlations, rather than the time at which correlations first become statistically significant, thereby establishing a conservative estimate of the interval over which correlations between neural activity and behavior can be detected.

**Figure 6.**
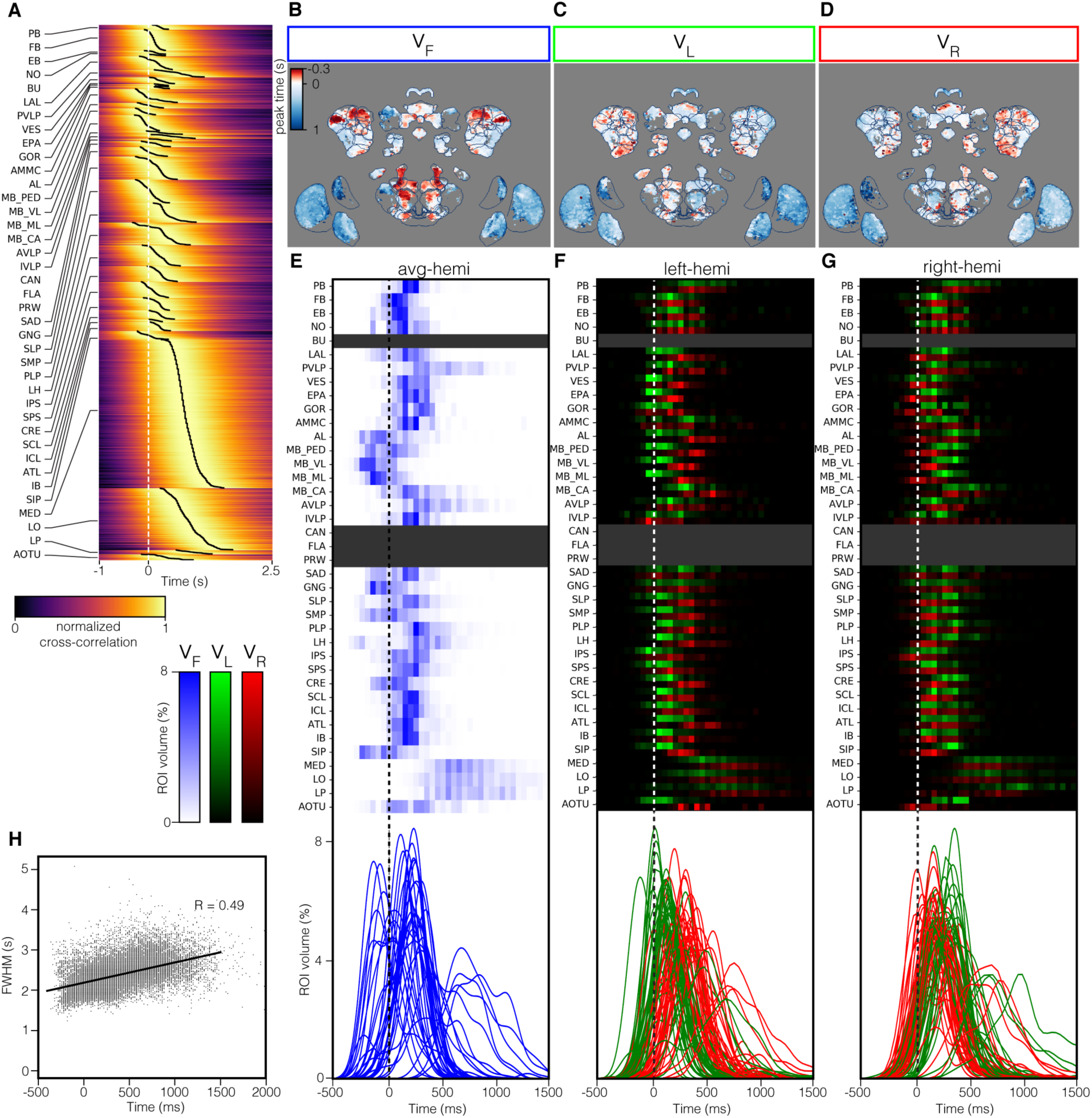
The temporal relationship between neural activity and behavior varies across brain regions. (A) Cross-correlation filters of supervoxel neural activity and forward velocity (n = 9 flies). Filters are first sorted by anatomical region, then peak time. Peak times are marked by the black traces. Filters are an average of both hemispheres., and have been deconvolved based on measurements of the impulse response of GCaMP6f kinetics (see Methods). (B) Maximum intensity projection. Colors represent time of peak cross-correlation between neural activity and forward velocity. Only filters that pass a significance threshold are shown (see methods) (n = 9 flies). (C) Same, but for left rotational velocity. (D) Same, but for right rotational velocity. (E) Histograms of peak cross-correlation of supervoxel neural activity and forward velocity. For each anatomical region, the percent of the volume that peaks at a given time is plotted. Below, kernel density estimates from each region are overlayed. (F) As in (E), but left and right rotational velocities for the left-hemisphere. (G) As in (E), but left and right rotational velocities for the right-hemisphere. (H) Relationship between forward velocity filter peak time and full-width-at-half-max. Each dot is a supervoxel. Black line is a linear regression.

To capture the temporal profiles of each brain region, we plotted a histogram of peak correlation times for each supervoxel within a region (Figure 6E-G). For changes in forward velocity, the earliest peak correlations appeared in olfactory sensory areas, namely the Antennal Lobe (AL) and Lateral Horn (LH), raising the possibility that these movements were initiated by incidental odors encountered by the fly inside the microscope (Figure 6E). Coincident with this, several subregions of the mushroom body are also highly correlated with future movements. Soon afterward, specific areas associated with extensive descending neuron innervation, such as the Gnathal Ganglion (GNG) and the Saddle (SAD) also display peak correlations with future changes in forward velocity, correlations that can persist until after the change in velocity has occurred. The Central Complex (including the Protocerebral Bridge (PB), the Fan-shaped Body (FB), the Ellipsoid Body (EB) and the Noduli (NO)) also include subregions that have peak correlations that span the initiation of movement, but also persist after the correlated change in forward velocity. Finally, peak correlations in sensory areas like the visual system (including the Medulla (MED) and the Lobula (LO), as well as the Lobula Plate (LP) emerged only after the change in velocity, and persisted in some cases for more than a second. Thus, changes in forward velocity are correlated with a characteristic sequence of changes in neural activity distributed widely across the brain, spanning sensory, motor, and navigation centers.

A different temporal sequence was observed when neural activity in each supervoxel was correlated with angular velocity (Figure 6F, G). In this case, while neural activity in olfactory regions as well as the mushroom body continued to correlate with future changes in angular velocity, a set of motor areas that partially overlapped with that associated with forward velocity but was nonetheless distinct, including the Inferior Posterior Slope (IPS) the Gorget (GOR), and the Vest (VES), were also correlated with future movements. Correlations between angular velocity and supervoxels in navigation centers, namely the Protocerebral Bridge (PB), the Fan-shaped Body (FB), the Ellipsoid Body (EB) and the Noduli (NO), lagged changes in velocity, consistent with the fact that these regions can integrate angular velocity signals to represent heading (Green and Maimon, 2018; Kim et al., 2017; Shiozaki et al., 2020). Signals in the optic lobes, the Medulla (MED), the Lobula (LO) and the Lobula Plate (LP), substantially lagged the correlated change in angular velocity. Finally, across nearly all brain regions, the time of peak correlation with angular velocity emerged first in the ipsilateral hemisphere relative to the contralateral hemisphere. Thus, the antagonistic relationship we observed between hemispheres using our linear modeling (Figure 5) appears to reflect an anticipatory increase in neural activity within specific regions of the ipsilateral hemisphere, followed by delayed, selective decrease of neural activity in the contralateral hemisphere. Finally, we note that maps of the correlations between neural activity and acceleration and deceleration show nearly identical temporal patterns to those seen using velocity (Figure S6).

### Individual brain regions display spatiotemporally structured patterns of engagement

Given the observation that individual brain regions contained topographic maps of behavioral variables, we next examined whether the distribution of peak correlation times we observed within a region might reflect spatiotemporal trajectories within each topographic map (Figure 7; Figure S7). These analyses revealed that even within each brain region, the temporal order of neural engagement was highly organized, and aligned with both behavioral and anatomical features. For example, the Lateral Horn (LH) contains a dorsal region that was more correlated with forward velocity, and a ventral region that was more associated with angular velocity (Figure 3). For forward velocity, this functional division corresponds to a temporal separation, with the dorsal region being correlated with behavior before the ventral region. Conversely, for angular velocity, this functional division is erased: on the ipsilateral side, most of the Lateral Horn (LH) displayed early correlations with the change in velocity, while on the contralateral side, peak correlation times gradually progressed along the medial-lateral axis. Indeed, such temporal gradients were widespread across brain regions, across different axes, and correlated with either forward or angular velocity, being apparent in the Inferior Posterior Slope (IPS), Superior Medial Protocerebrum (SMP), the Superior Intermediate Protocerebrum (SIP), and the Fan-Shaped Body (FB). In other brain regions, such as the Lateral Accessory Lobe (LAL), the Mushroom Body Medial Lobe (MB-ML), and the Posterior Lateral Protocerebrum (PLP), the timing of peak correlations with forward velocity were uniformly distributed across the structure, while the timing of peak correlations with angular velocity displayed the characteristic temporal order of ipsilateral signals preceding contralateral signals. Finally, in the optic lobe, where all of the correlations with behavior presumably reflect feedback entering the visual system from the central brain, the temporal order of activation captured the ipsi-leading, contra-lagging pattern for angular velocity, and mapped onto distinct “input” layers.

**Figure 7.**
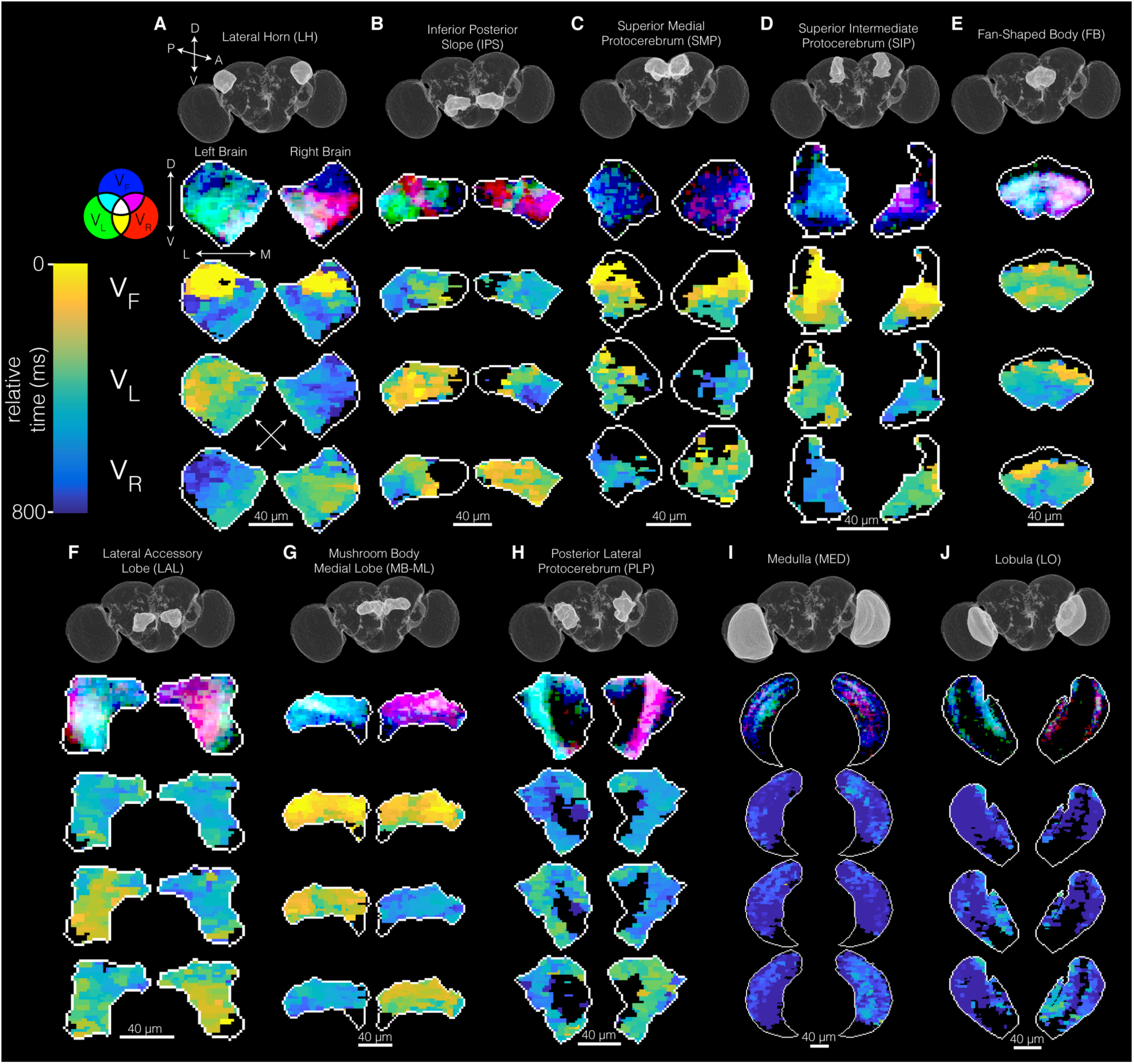
Topographic maps of peak correlation times are spatially structured within individual brain regions. (A). Lateral Horn (LH). (B) Inferior Posterior Slope (IPS). (C) Superior Medial Protocerebrum (SMP). (D) Superior Intermediate Protocerebrum (SIP). (E) Fan-shaped Body (FB). (F) Lateral Accessory Lobe (LAL). (G) Mushroom Body Medial Lobe (MB-ML). (H) Posterior Lateral Protocerebrum (PLP). (I) Lobula (LO). (J). Medulla (ME). Each column contains the correlation maps describing the categorial voxel assignments (from Figure 3), as well as the topographic maps of relative peak correlation time (where the earliest peak correlation time relative to behavior is set to 0) for Vf, Vl and Vr.

Taken together, these results demonstrate that the temporal relationships between neural signals in each brain region and changes in specific behavioral velocities is exceptionally rich, and reflect multiple levels of functional and anatomical organization. Strikingly, temporal gradients of correlation that presumably reflect wave-like sequences of changes in neural activity that propagate across specific regions are a common organizational principle in this system.

## Discussion

### Summary

Here we describe a novel two-photon imaging approach to extract neural signals across the entire *Drosophila* brain as the animal behaves, and via volumetric registration, quantitatively compare signals across brain regions and individuals (Figure 1). We find that neural activity containing information about locomotion is widespread across the brain, extending well beyond regions commonly associated with motor control (Figure 2). We observe striking topographic order within individual brain regions, such that neurons with distinct selectivity for behavioral features are highly compartmentalized (Figure 3). Remarkably, these signals account for the dominant dimensions of neural activity in the brain (Figure 4). These signals include a nearly brain-wide state-change associated with the transition between moving and not moving, as well as localized signals that contain information specific to the forward or rotational velocity of the fly (Figure 5). We find that the temporal relationship between neural activity and behavior evolves across the brain: activity in some regions precedes changes in locomotor velocity by 300 ms; in others, changes in neural activity are contemporaneous with behavior, while in others, neural activity lags behavioral changes by more than a second (Figure 6). Strikingly, neural signals associated with changes in angular velocity are anti-symmetric across hemispheres, such that an increase in activity in several brain regions on the ipsilateral side precedes a mirrored decrease in activity in the same regions on the contralateral side (Figure 5, 6). Within many individual brain regions we observe temporal gradients of neural activity that sometimes respect functional compartments, but more often sequentially engage neurons with different behavioral selectivity (Figure 7). Taken together, these studies identify a brain-wide spatiotemporal topography of walking, in which distinct networks of neurons are engaged by specific behavior maneuvers and follow stereotyped temporal trajectories (Figure 8).

**Figure 8.**
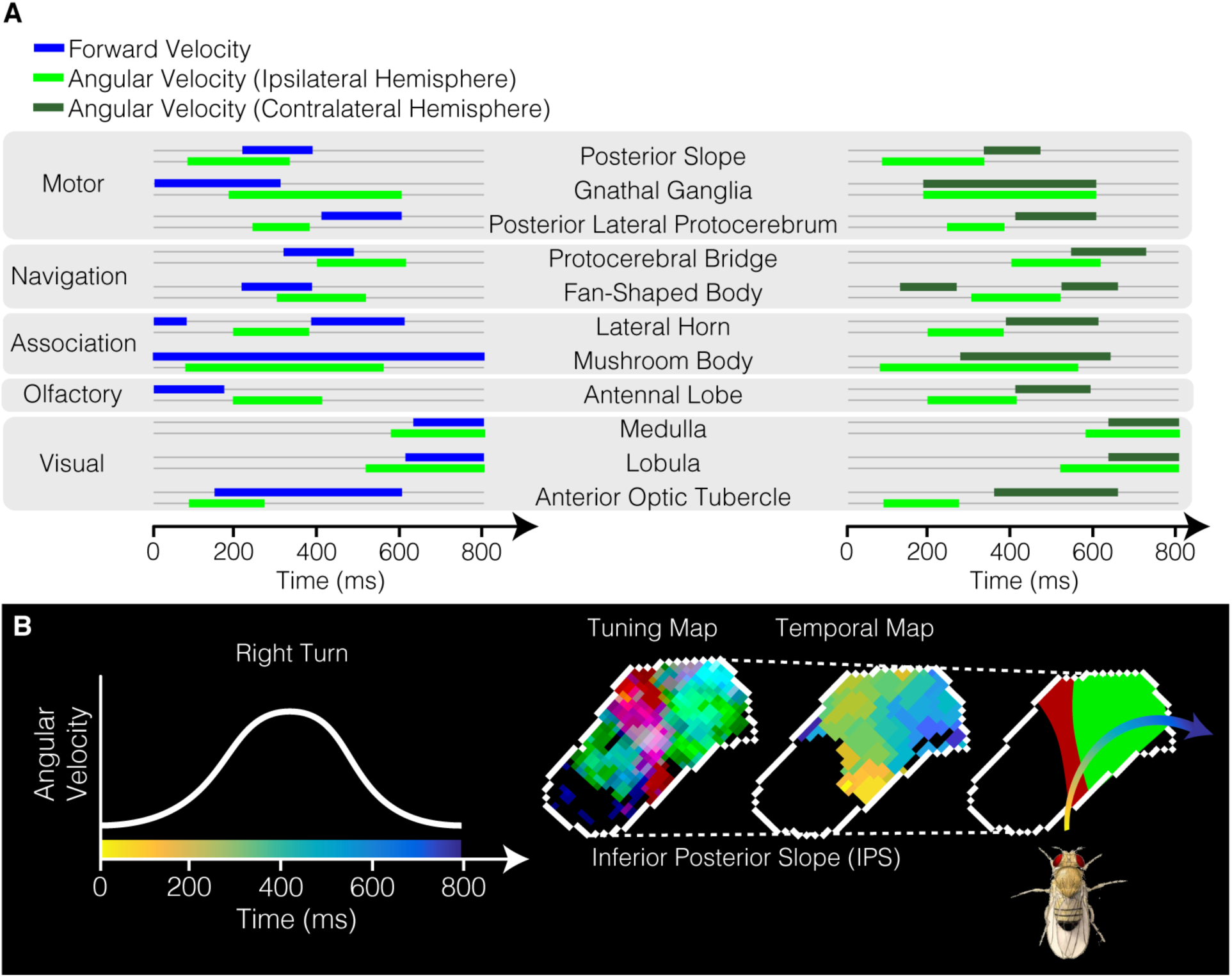
Locomotion related signals are temporally ordered within and across brain regions. (A) Abstract representation of categorized brain regions activating and deactivating along a temporal sequence. Comparing forward and angular velocity, as well as ipsilateral and contralateral hemispheres for angular velocity. (B) Abstract representation of how a motor region with compartmentalized velocity tuning combined with a temporal gradient of neural activity could embed a stereotyped behavioral maneuver, such as a turn.

### A unified functional brain space enables quantitative comparisons across individuals and conditions

Spatial registration of neural data in *Drosophila* has enabled genetically labeled neurons in different animals to be compared in the same space, and has allowed functional data to be aligned with established brain atlases (Bates et al., 2020; Bogovic et al., 2020; Costa et al., 2016; Ito et al., 2014; Mann et al., 2017; Mann et al., 2021; Münch et al., 2021; Pacheco et al., 2021; Peng et al., 2011). Here we describe a generalizable, fully automated pipeline for functional brain registration in behaving animals, spanning virtually the entire brain. We demonstrate how spatial registration in the context of a brain whose wiring is highly stereotyped between individuals can be leveraged to increase the SNR of each voxel. Moreover, we show that registration enables modeling of neural and behavioral data across individuals without compromising spatial or temporal resolution. Most significantly, the capacity to routinely register functional signals from many individual brains into a common space enables future quantitative comparisons across experimental conditions and laboratories. Using this approach, future work could compare functional maps of different sensory modalities, behaviors, and neural computations, as well as conditions that would be impossible to implement in a single animal, such as silencing or activating many different genetically targeted populations of neurons, or exploring the effects of particular genes on functional maps. Finally, registering these functional data with fly connectomes will greatly increase opportunities for modeling how neural signals are transformed across the brain (Scheffer et al., 2020; Turner et al., 2021; Zheng et al., 2018).

### Locomotor Signals are Widespread and Highly Structured

Measurements and perturbations of genetically identified cell types have provided fundamental insights into a wide diversity of neural processes in the fruit fly. At the same time, how these relatively compact circuit computations might be embedded within larger neural networks that could be engaged during behavior has remained unclear. Indeed, while data in vertebrates demonstrates that behavioral signals are prominent in many different brain regions, whether such a distributed representation of locomotion might exist in the fly was unknown. Our data demonstrate that locomotor signals can be detected in approximately 40% of the brain volume, and across almost every neuropil (Figure 2). Some of this signal can be attributed to a global change in activity that is associated with the onset of movement (Figure 4). However, much of this signal is selective to specific behavioral variables and brain regions (Figure 5). This selectivity is spatially structured, even within individual brain regions, such that neural activity signals with different selectivities are often topographically segregated at the mesoscale, well above the spatial resolution of our imaging approach (Figure 3). Given the scale of these motor maps, this observation is consistent with the notion that neurons that have similar tuning with respect to locomotor behavior are physically grouped.

These distributed locomotor signals likely play diverse computational roles in different brain regions. For example, modulation of visual circuits by locomotion-evoked changes in octopamine increase the gain of visual processing to capture the more rapid changes in visual scene statistics caused by movement (Suver et al., 2012). In addition, locomotion can provide a predictive signal that can incorporate the dynamics of locomotor behavior to modulate motion processing (Cruz et al., 2021; Fujiwara et al., 2017; Kim et al., 2015; Kim et al., 2017; Strother et al., 2017). Consistent with these previous observations, our data reveal extensive locomotor signals in the visual neuropils, including the Medulla, Lobula, and Lobula Plate, and are often restricted to specific processing layers (Figure 5). We note that we were unlikely to directly detect rapid efference copy signals that hyperpolarize specific visual interneurons given that we were imaging GCaMP6f, which favors depolarizing signals (Chen et al., 2013; Fujiwara et al., 2017; Kim et al., 2015; Kim et al., 2017).

Beyond early visual processing, behavior signals have been previously observed in higher-order circuits. In particular, the mushroom body, responsible for learning and memory in *Drosophila*, encodes behavior as a dopaminergic signal to coordinate synaptic plasticity and reward (Cohn et al., 2015; Zolin et al., 2021). In addition, navigation circuits in the ellipsoid body rely on behavior evoked signals to coordinate remapping of the heading-direction network (Fisher et al., 2019). Moreover, the fan-shaped body, another region engaged with sensory integration and navigation, shows behavior-based gating of visual responses (Weir and Dickinson, 2015). Our data extend these observations to many other regions in the central brain that have had only limited functional characterization, opening new avenues for understanding the interactions between behavior and other circuit computations. Finally, the breadth and diversity of the locomotor signals we describe in the fly closely parallel analogous observations in a variety of contexts and brain regions in other animals (Kaplan and Zimmer, 2020).

### Different locomotor movements engage distinct spatiotemporal neural networks

Locomotor behavior reflects complex, time-evolving changes in specific velocities that are structured over hundreds of milliseconds (Berman et al., 2014; Branson et al., 2009; Chun et al., 2021; DeAngelis et al., 2019; Katsov et al., 2017; Kain et al., 2013; Mendes et al., 2013; Strauss and Heisenberg, 1990). Our data demonstrate that changes in the instantaneous forward and angular velocities of the animal engage distinct neural networks distributed over multiple brain regions, and follow stereotyped temporal sequences of coordinated activity (Figure 8). While the network that is engaged by changes in forward velocity is bilaterally symmetric, the networks engaged in changes in angular velocity are anti-symmetric across the midline, revealing that activation of specific areas in one hemisphere is paired with a subsequent, targeted reduction in signal in the same regions in the other hemisphere. As analogous antisymmetric networks for steering control have long been known from work in vertebrate spinal cord (Deliagina et al., 2000; Fagerstedt et al., 2001; Grillner et al., 2007), our data demonstrate that this organizational principle extends across the brain.

Focusing on neural signals that anticipate and are contemporaneous with changes in behavioral velocities reveals engagement of regions that contain substantial innervation by descending neurons, a key bottleneck in motor control, as well as other regions whose relationship to motor control is unknown. Thus, understanding how temporal patterns of descending neuron recruitment and dismissal emerge will likely require relating these patterns to the extensive networks in which they are embedded.

The fine structure of our spatiotemporal maps suggest a framework for how neurons with different feature selectivities can be sequentially engaged during a movement trajectory (Figure 8). In particular, while neurons with similar selectivities are spatially clustered, the temporal precession of neural activity is often spatially graded, and can cross more than one cluster. This suggests that different pools of neurons are recruited in a specific temporal order that relates to the evolving pattern of movement. Such a dynamical systems perspective parallels the conceptual framework that describes forelimb movement in primate motor cortex (Churchland et al., 2012; Shenoy et al., 2013). In this view, populations of neurons are activated in a temporal pattern that describes the structure of the limb trajectory, but no spatial order has been detected within these populations. Our data raise the possibility that a conceptually similar principle underpins walking behavior in the fly, but that in this compact nervous system, a mesoscale spatial order has been imposed on the dynamical system. Given the powerful tools available in the fly for targeted measurements and perturbations, future work should define the causal structure of this dynamical system with single cell precision.

## Author Contributions

L.E.B. and T.R.C. conceived the project. L.E.B. collected experiments and analyzed the data.

A.B.B. registered neural data with Synthmorph. S.D. provided modeling direction and feedback.

L.E.B. and T.R.C. wrote the manuscript. T.R.C. advised throughout the project.

## Acknowledgements

We would like to thank the Dickinson lab for providing us with treadmill balls, as well as the Maimon lab for direction on building our fly mounting apparatus. We would also like to thank members of the Clandinin Lab, as well as Diego Pacheco, Osama Ahmed, and Salil Bidaye for feedback on the manuscript. This work was supported by a National Science Foundation Graduate Research Fellowship (LEB), as well as grants from the NIH (R01EY022628, 5U19NS104655 (TRC and SD), 3R01NS11006003 (TRC) and the Stanford Vision Core Grant 5P30EY02687704 (TRC)) and the Simons Foundation (TRC).

## Supplemental Figures

**Supplemental Figure 1.**
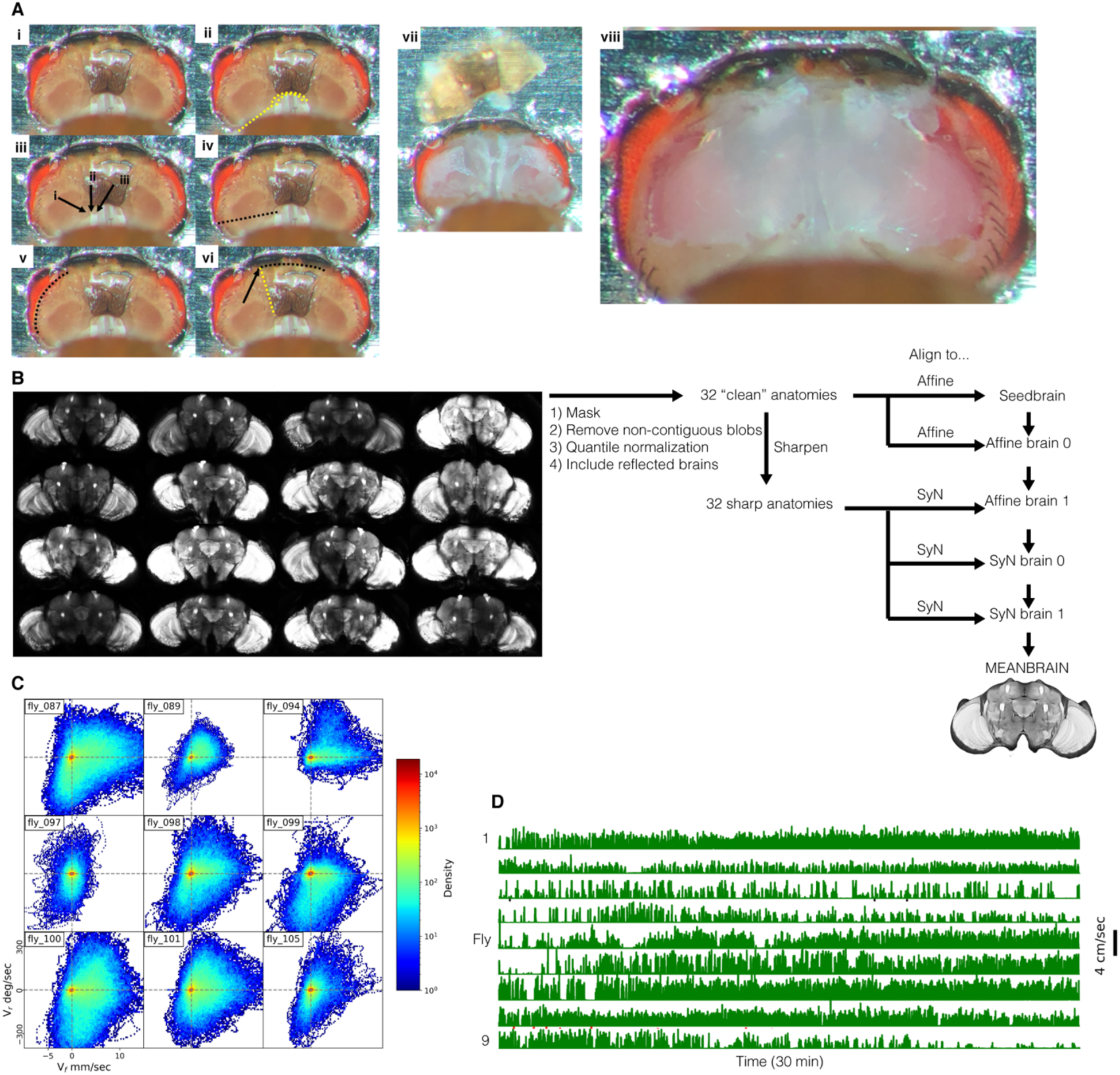
Whole-brain imaging and data alignment in walking *Drosophila*, related to Figure 1. (A) Overview of the sequence of dissections needed to expose the entire brain. (i) Each fly was glued into a metal shim. (ii) Placement of first cuts with dissection needle. (iii) Three cuticle boundaries near the neck. (iv) Cut along the black line. (v) Cut along the eye border. (vi) Yellow line denotes a strong cuticular structure. Continue cutting along the black line, breaking through yellow line. (vii) Remove cuticle. (viii) Remove trachea and fat. (B) Meanbrain creation. Slices through 16 anatomical volumes (including 9 brains from animals that were used for functional imaging, as well as 7 additional animals) were used to create the meanbrain. (C) 2D histograms of Vf, Vr and Vl for the nine flies in this study. (D) Temporal traces of walking behavior derived from ball movement across the entire 30 minute session for each fly.

**Supplemental Figure 2.**
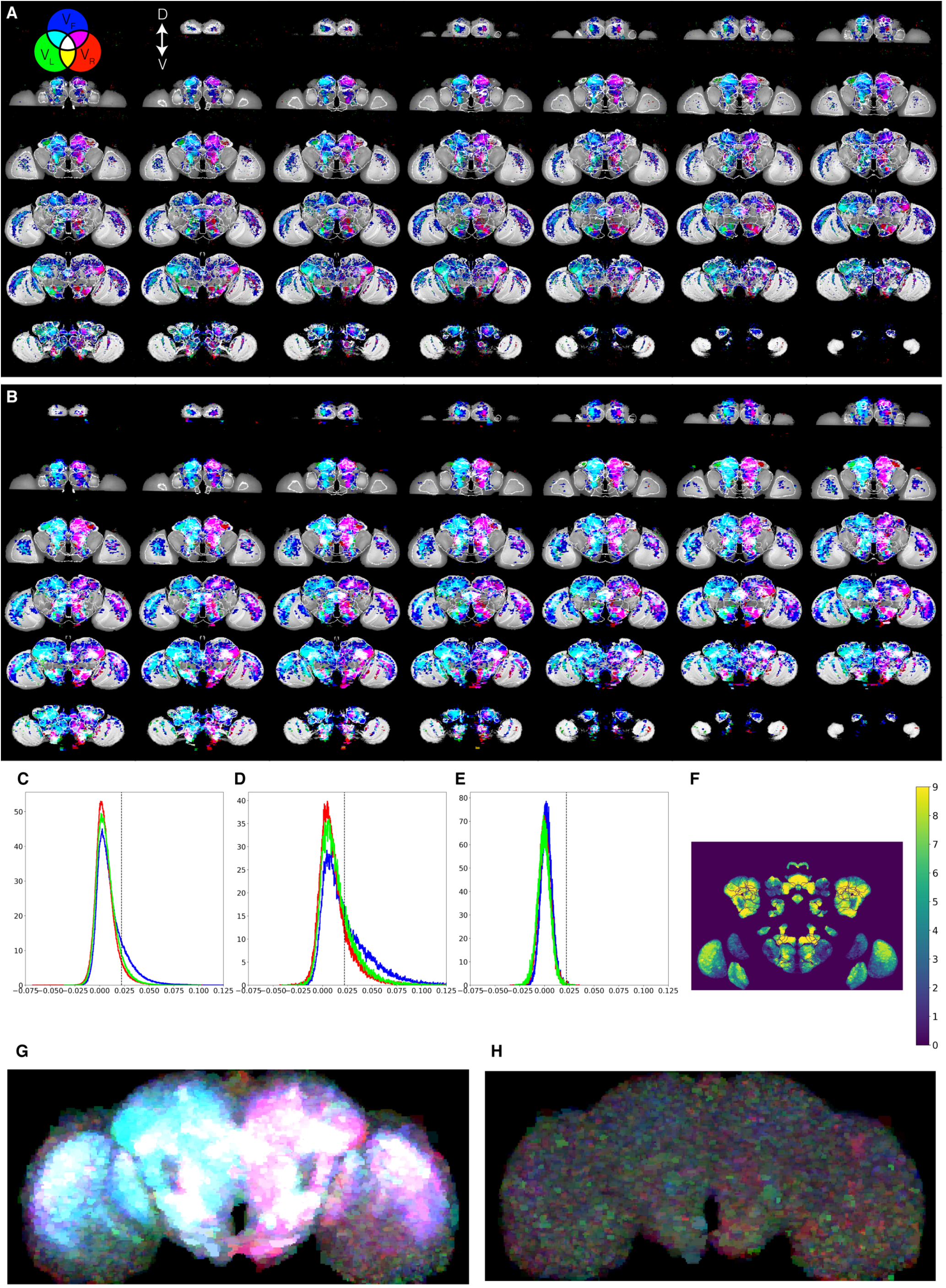
Brainwide correlations measured with single voxels versus supervoxels using both GcAMP and tdTomato to control for brain motion, related to Figure 2. (A) Slices through the meanbrain overlaid with velocity component correlations to single voxels and anatomical ROI boundaries. Slices are arranged from anterior to posterior. See Figure 2. (B) As in (A) but with supervoxels instead of single voxels. (C) Distribution of single voxel correlations with three velocity components. Dashed line indicates the significance threshold used in Figure 2. (D) As in (C), but with supervoxels. Note increased correlations. (E) As in (D), but calculating the correlations using the tdTomato signal (structural marker) instead of GCaMP6f (neural activity). (F) Brain-wide map of the number of flies that had a correlation with velocity above an r-value of 0.05. (G) Maximum projection through the slices shown in (A). (H) As in (G), but using the correlations calculated with the tdtomato signal. The absence of signal in this Figure indicates that motion artifacts were undetectably small.

**Supplemental Figure 3.**
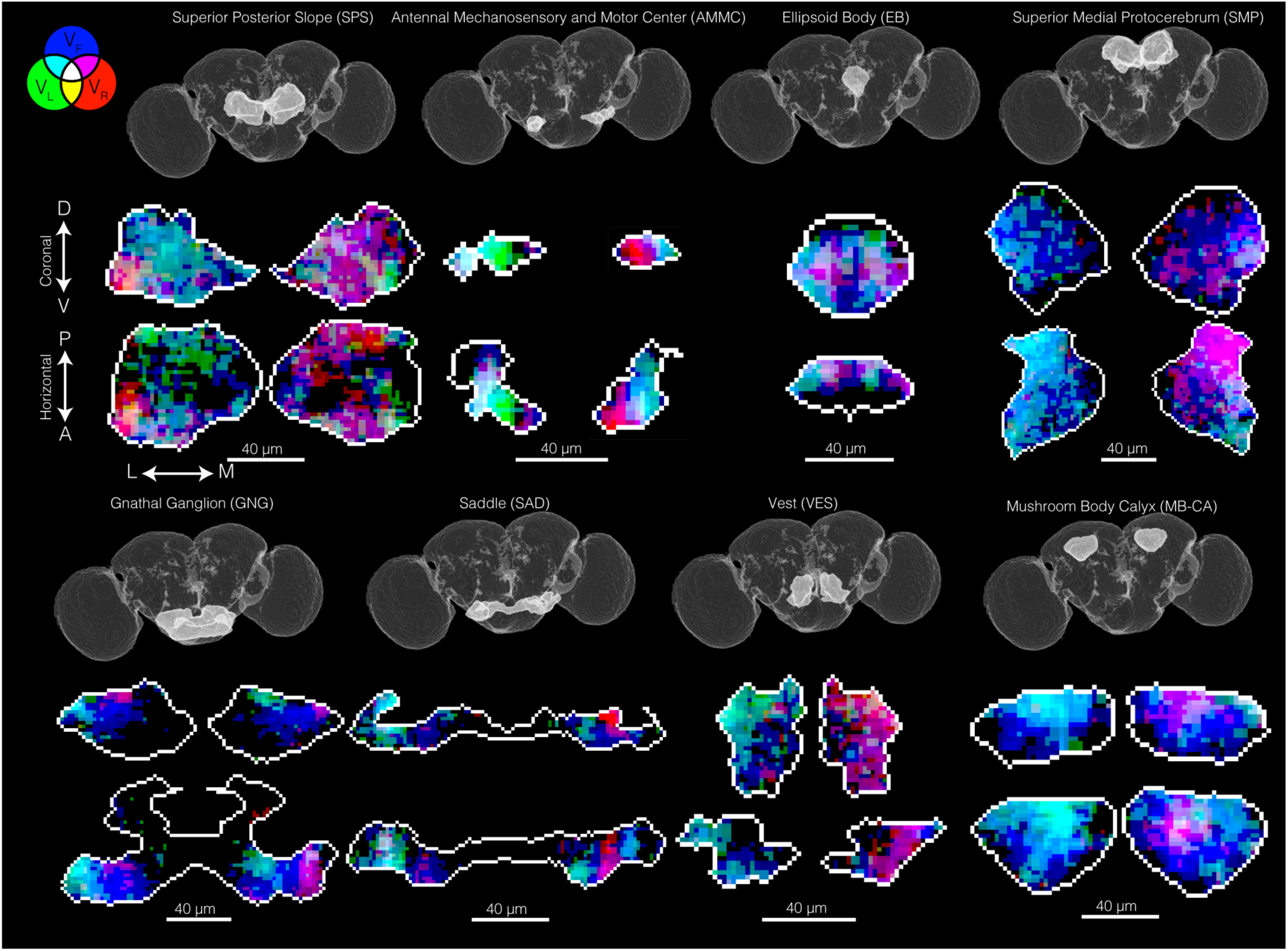
Topographic maps of neural tuning to behavioral velocities in specific brain regions, related to Figure 3. Slices of correlation maps through the Superior Posterior Slope (SPS), the Antennal Mechanosensory and Motor Center (AMMC), the Ellipsoid Body (EB), the Superior Medial Protocerebrum (SMP), the Gnathal Ganglion (GNG), the Saddle (SAD), the Vest (VES), and the Mushroom Body Calyx (MB-CA).

**Supplemental Figure 4.**
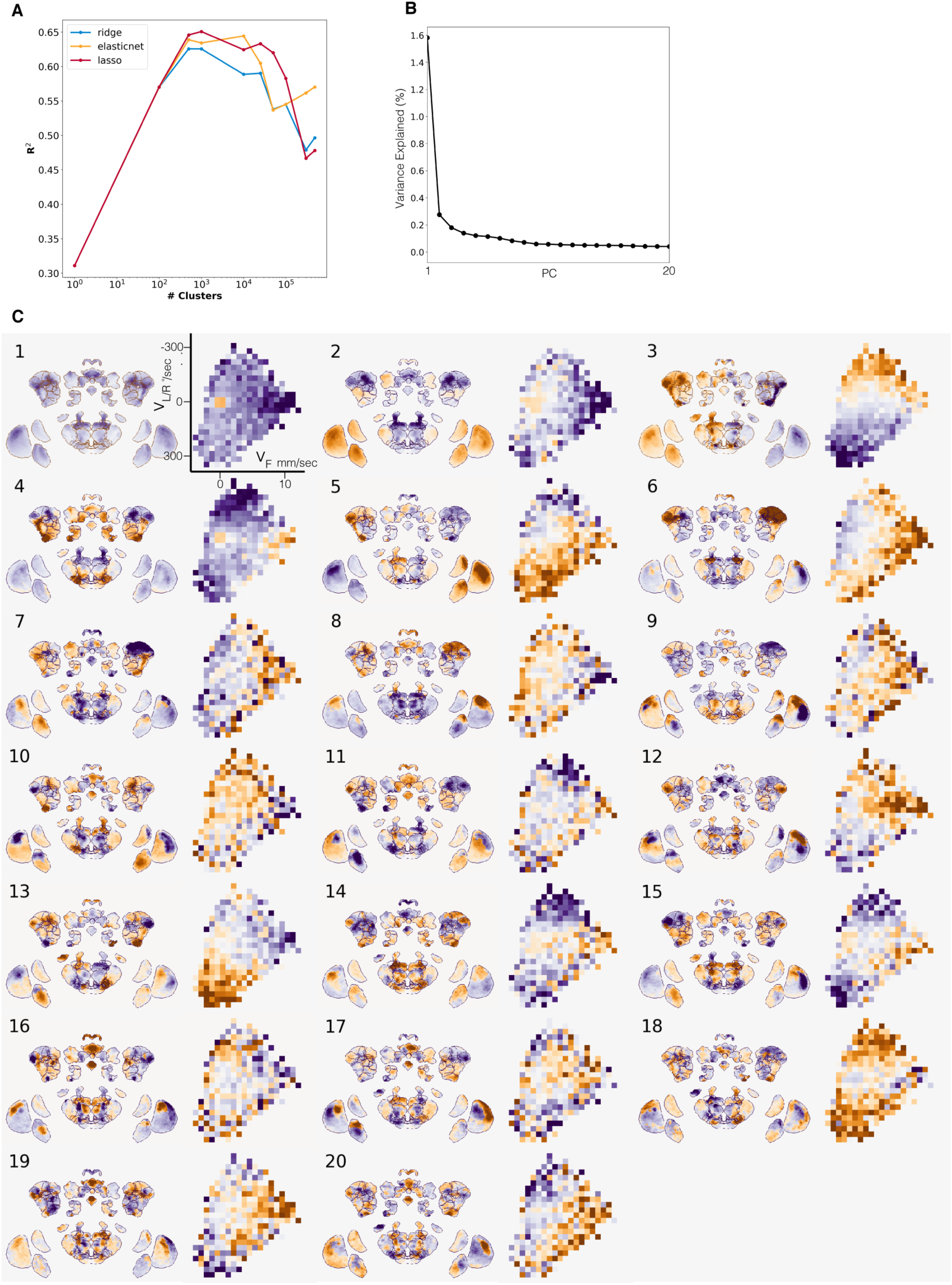
Many dimensions of neural activity are predictive of velocity space, related to Figure 4. (A) Voxels were agglomerated into supervoxels before performing PCA. To determine the optimum number of supervoxels per slice, we swept a range of supervoxel sizes, and fit linear models that predict the forward velocity of the fly. Too few or too many supervoxels reduced accuracy. Three regularization methods were used. (B) Neural variance explained by the first 20 principal components. (C) The first 20 principal components viewed on the brain and in 2D behavior space, as in Figure 4.

**Supplemental Figure 5.**
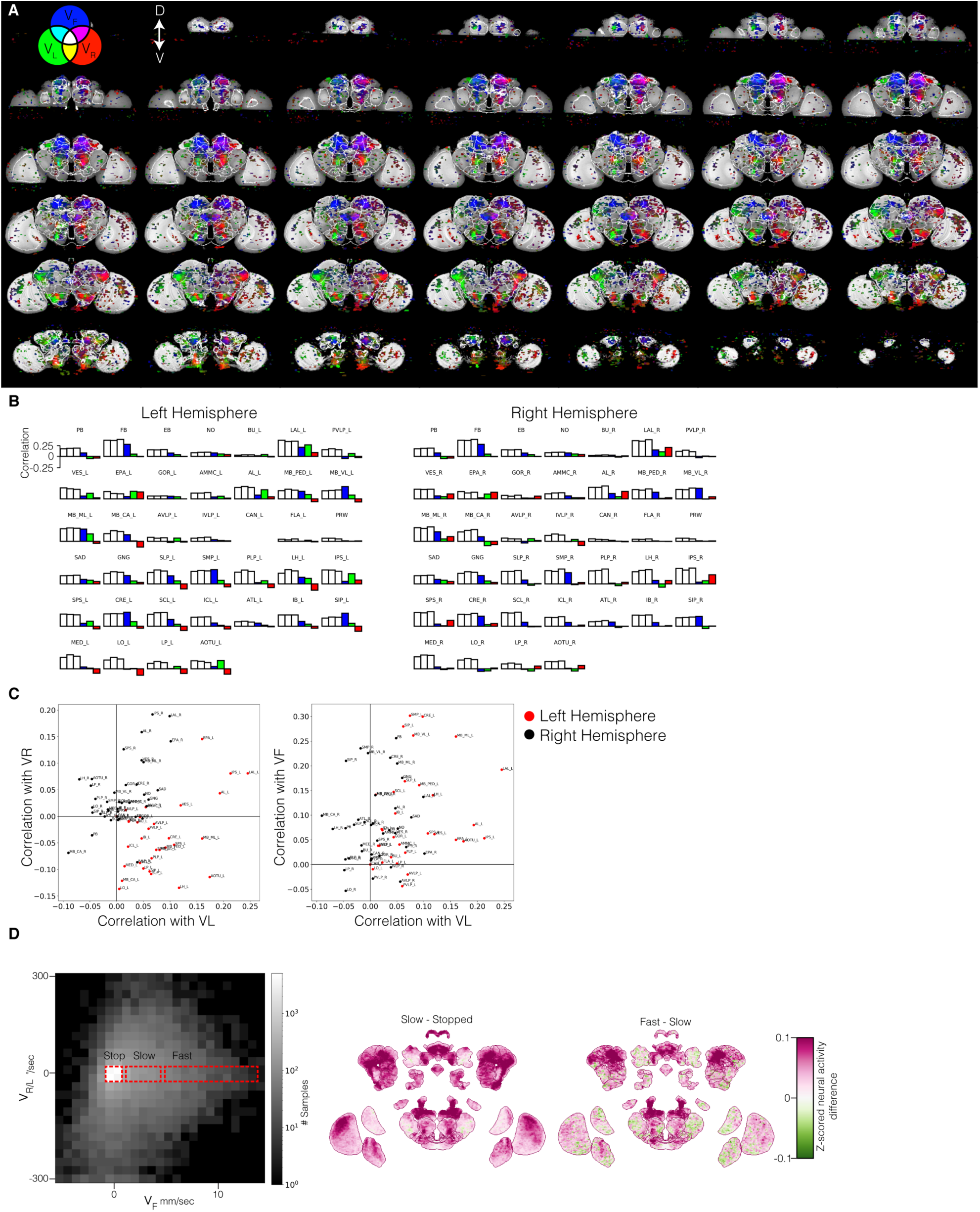
Topographic inspection of velocity demixing results, related to Figure 5. (A) Slices through the meanbrain, overlaid with demixed velocity component R^2^ to supervoxels and anatomical ROI boundaries. Slices are arranged from anterior to posterior. See Figure 5. (B) As in Figure 5C, but reporting average correlations across anatomically defined regions. White bars were below the movement threshold; color bars denote velocity components above the movement threshold. (C) Above movement threshold correlations with each anatomically defined region and velocity component. (D) Extension of Figure 5F, but showing alternative 2D behavior defined comparisons. Comparing the difference in neural activity across the brain for a fly stopped versus moving slowly forward, and a fly moving slowly forward versus quickly forward.

**Supplemental Figure 6.**
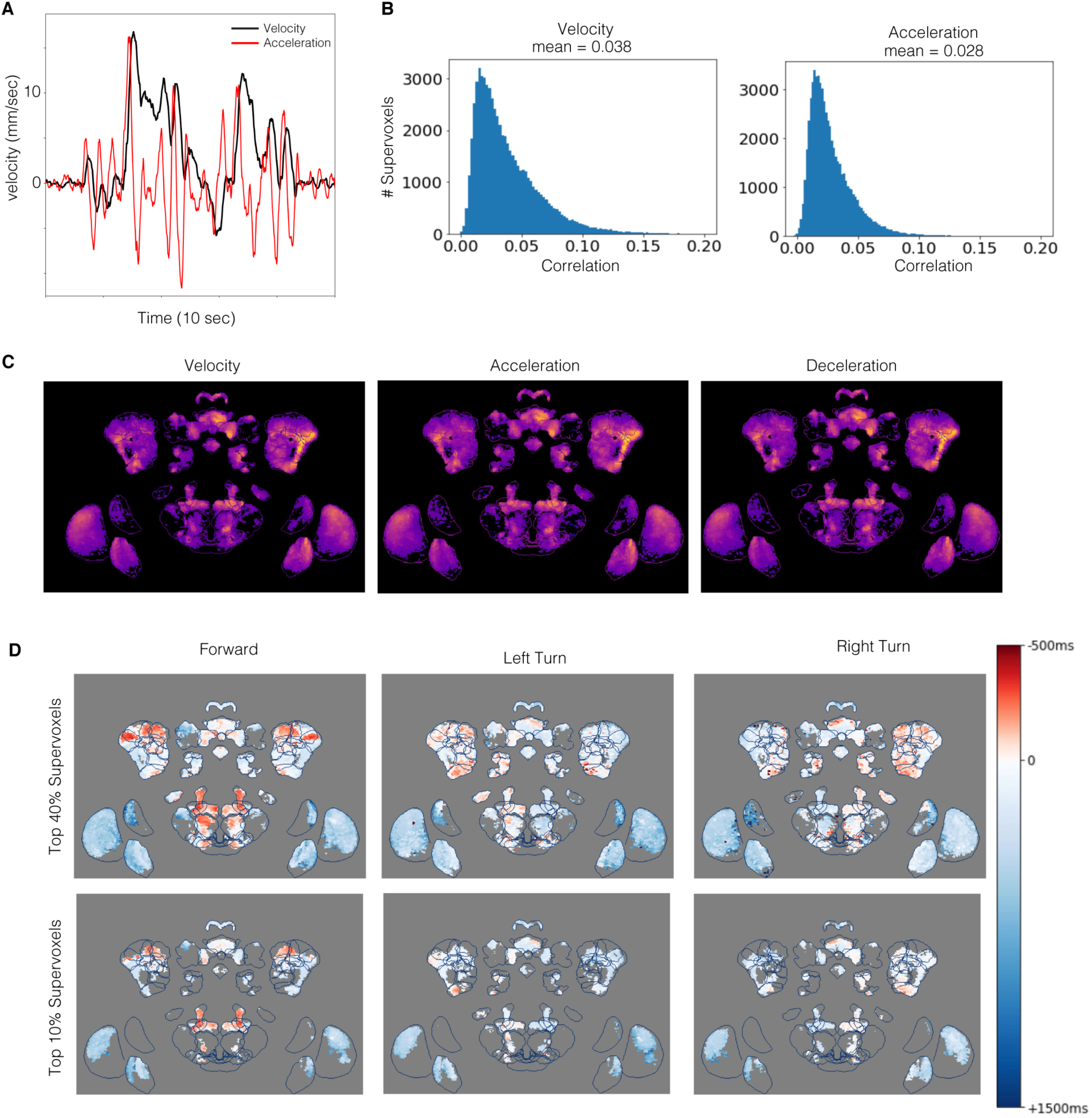
Temporal relationship between neural activity, velocity, and acceleration, related to Figure 6. (A) Trace of forward velocity and acceleration for a 10 second window of behavior. (B) Histograms of instantaneous correlation between neural activity of supervoxels and forward velocity or acceleration. (C) Comparing peak cross-correlation values for velocity, acceleration, and deceleration for a right turn. (D) Map of peak cross-correlation time between neural activity and velocity components. Comparing two thresholds for significant voxels.

**Supplemental Figure 7.**
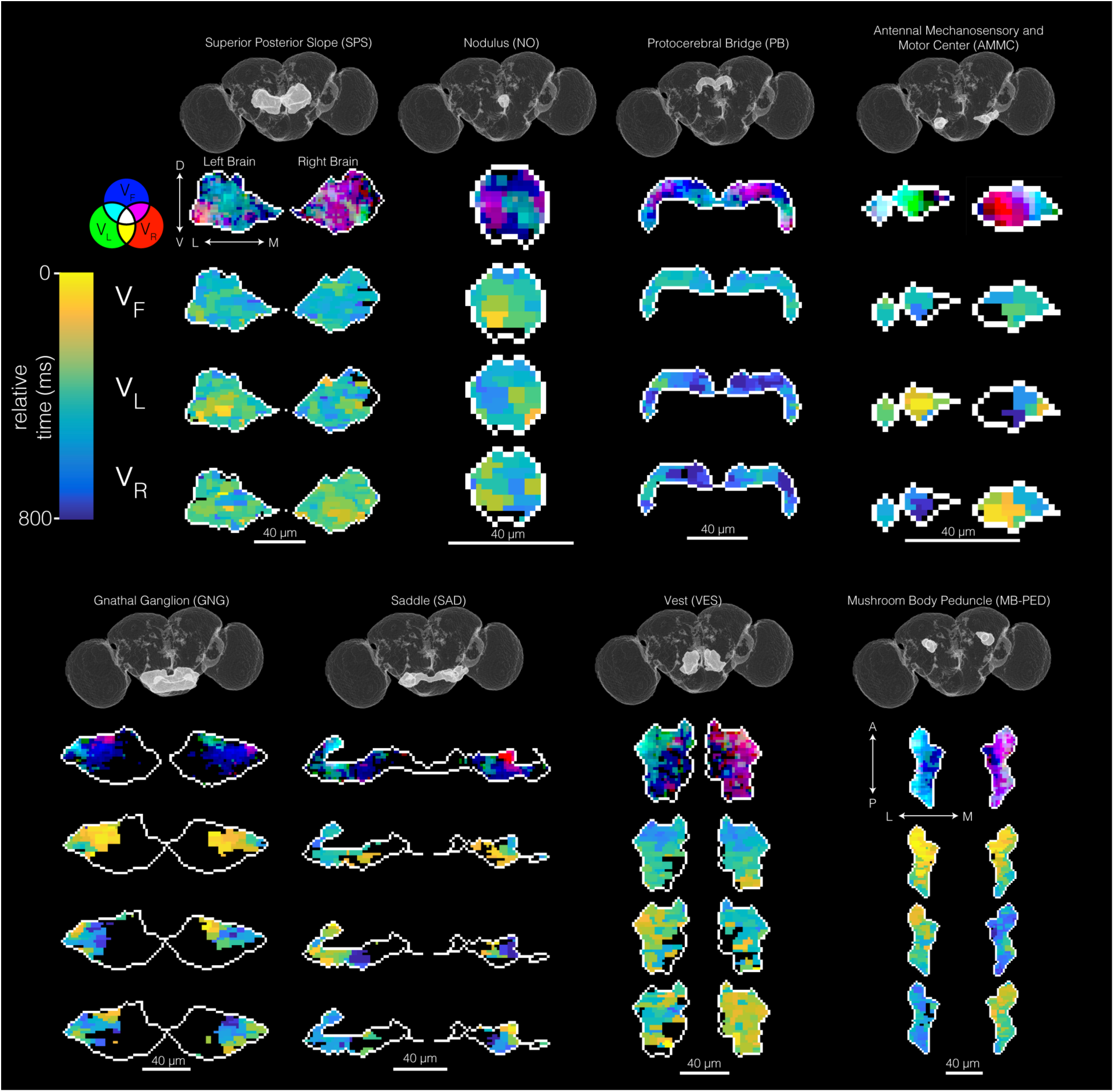
Topographic maps of peak correlation times are spatially structured within individual brain regions, related to Figure 7. Slices through individual brain regions showing velocity component tuning and peak cross-correlation. Regions are Superior Posterior Slope (SPS), Nodulus (NO), Protocerebral Bridge (PB), Antennal Mechanosensory and Motor Center (AMMC), Gnathal Ganglion (GNG), Saddle (SAD), Vest (VES), Mushroom Body Peduncle (MB-PED).

## STAR Methods

### RESOURCE AVAILABILITY

#### Lead contact

Further information and requests for resources should be directed to the lead contact, Thomas R. Clandinin.

#### Materials availability

This study did not generate new unique reagents.

#### Data and code availability

- Microscopy (structural and functional neural recordings) and behavioral data have been deposited at Dryad and are publicly available as of the date of publication. DOIs are listed in the key resources table.
- All original code has been deposited at Github and is publicly available as of the date of publication. DOIs are listed in the key resources table.
- Any additional information required to reanalyze the data reported in this paper is available from the lead contact upon request.

### EXPERIMENTAL MODEL AND SUBJECT DETAILS

*Drosophila melanogaster* were of the genotype *w+/w+;UAS-myr::tdTomato/UAS-GCaMP6f;nSyb-Gal4/+*. Flies were raised on molasses medium at 25 °C with a 12/12-h light/dark cycle. Flies were housed in mixed male/female vials of 10-20 individuals. 3-4 days post-eclosion females were used for imaging.

### METHOD DETAILS

#### Mounting and Dissection

Each fly was anesthetized on a chilled Peltier plate with a thermally coupled custom holder. Each immobilized fly was carefully fitted into a custom mount consisting of 3D-printed plastic and a custom cut steel shim to tightly nestle the head and thorax. To fix the fly to the mount, UV-curable glue was placed and cured on the dorsal region of the face between the eyes, and on the thorax. A saline solution was added to the dish for dissection (103 mM NaCl, 3 mM KCl, 5 mM TES, 1 mM NaH_2_PO_4_, 4 mM MgCl_2_, 1.5 mM CaCl_2_, 10 mM trehalose, 10 mM glucose, 7 mM sucrose, and 26 mM NaHCO_3_). Using a tungsten needle the posterior head cuticle was carefully cut and removed to reveal the whole brain (Figure S1). Dissection forceps were used to remove fat and trachea.

#### Two-Photon Imaging

Flies were imaged using a resonant scanning Bruker Ultima IV system with a piezo drive and a Leica 20× HCX APO 1.0 NA water immersion objective lens. GCaMP6f and tdTomato were simultaneously excited with a Chameleon Vision II femtosecond laser (Coherent) at 920 nm. A 525/50 nm filter and a 550/50 nm filter were used to collect signals from GCaMP6f and tdTomato. Photons were detected simultaneously using two GaAsP-type photomultiplier tubes. The exposed fly brain was perfused with carbogen-bubbled (95% O_2_, 5% CO_2_) saline solution (same as above) heated to 30°C with an in-line heater. For the 30 min functional scan, volumes were collected at a resolution of 2.6 × 2.6 × 5 µm (256 voxels x 128 voxels x 49 slices, XYZ), resulting in an approximate volume rate of 1.8Hz. Scans were bidirectional along the X axis. For the immediately subsequent anatomical scan, spatial dimensions were adjusted to 0.6 × 0.6 × 1µm (1024 voxels x 512 voxels x 241 slices, XYZ), and 100 volumes were collected. In this orientation, all regions of the brain were visible, except for the laminas in each optic lobe, which are occluded by the eye, and a portion of the Gnathal Ganglion, which is occluded by the esophagus.

#### Behavior Tracking

During imaging, the head-fixed fly performed spontaneous bouts of walking on a painted, air-suspended foam ball (9 mm diameter, LAST-A-FOAM FR4615). The ball was imaged at 50 Hz with a Flea FL3-U3-13E4M-C sensor and Edmund Optics 100 mm C Series Fixed Focal Length Lens. An IR LED directed with optic fibers was used to illuminate the ball. Frames were processed using Fictrac to calculate the animal’s walking velocity (Moore et al., 2014). Before all subsequent analysis, forward and rotational velocities were smoothed with a Savitzky-Golay filter of window length 500 ms and a polynomial of order 3.

#### Data Preprocessing

Brain volumes were first motion-corrected using ANTs (Avants et al., 2009; Avants et al., 2011); the tdTomato channel was time-averaged across the 30 min recording and each tdTomato volume was warped (affine and non-linear) to the mean. Each volume’s warp parameters were then applied to the GCaMP6f channel. Then, each voxel was independently corrected for bleaching as well as other slow temporal trends by subtracting a temporally smoothed signal from the raw trace (smooth signal produced by gaussian filter of 2 minute sigma, truncated at 1 sigma). Finally, each voxel in the GCaMP6f recording was Z-scored. This preprocessing was all done on individual animals before volumetric alignment and concatenation.

#### Data Alignment

Each individual’s anatomical scan was created from 100 collected volumes of tdTomato signal. These 100 volumes were first averaged, then each volume was warped (affine and non-linear) to this mean using ANTs. These aligned volumes were then averaged, creating the final anatomical scan for the individual. A cross-individual mean anatomical brain was then produced to facilitate warping all data into a common space as follows. 16 anatomical scans, selected as the highest quality from a larger pool of approximately 30 brains, were first pre-processed with an intensity-based masking, removal of non-contiguous blobs, and quantile normalization to brighten overly-dark areas and darken overly-bright areas. Each brain was additionally mirrored across the Y-axis to double our effective data to 32 brains. These 32 brains were all aligned (affine) to a seed-brain chosen from the 32, and averaged to produce affine_0. The 32 brains were then aligned (affine) to affine_0, and averaged to produce affine_1. Next, the individual anatomical scans were sharpened (using unsharp masking), and aligned (affine and non-linear) to affine_1, and averaged to produced SyN_0. The last step was repeated two more times on the subsequent output brains to produce the final meanbrain. Finally, the meanbrain was affine aligned to the JFRC2018 female brain to put the brain in familiar coordinates as well as level the brain. With the mean brain in hand, we then aligned all our functional data into this space as follows. Each individual’s functional scan was affine aligned to their anatomical scan using the tdTomato channel and applying the transforms to the GCaMP6f channel. Then, each anatomical scan was aligned (affine and non-linear) to the meanbrain using the tdTomato channel, and the transforms were applied to the GCaMP6f channel. Finally, we warped an atlas of spatially defined anatomical brain regions into our meanbrain space by first warping the IBNWB brain (and accompanying atlas) into JRC 2018 female brain space, and then warping this into our meanbrain (affine and non-linear) (Ito 2014, Bogovic 2020). Warping JRC 2018 into our meanbrain space with ANTs was followed by SynthMorph to improve registration accuracy (Hoffmann 2020).

### QUANTIFICATION AND STATISTICAL ANALYSIS

#### Agglomerative Clustering Supervoxel Creation

Individual voxels were spatially aggregated into supervoxels to reduce the number of features for computational tractability and to boost SNR as follows. After functional GCaMP6f data was preprocessed and warped into the common meanbrain space, individual flies were temporally concatenated to create a single large “superfly” matrix of (x, y, z, t; 256, 128, 49, 30456) (Figure 1A). Then, supervoxels were merged independently for each z-slice via agglomerative clustering with Ward linkage and a connectivity constraint (only spatially neighboring voxels could be merged). The number of supervoxels per slice was determined by sweeping across a range of supervoxels (1, 100, 500, 1000, 2000, 5000, 10000) and for each fitting a linear model that takes the supervoxel neural signals and predicts behavior. Too few or too many supervoxels results in lower prediction accuracy, with a peak at 2000 supervoxels (Figure S4A).

#### Correlation Analysis (Voxel- and Supervoxel-Wise)

We started with the superfly with supervoxels (supervoxel, z, t; 2000, 49, 30456). The Pearson correlation was calculated for each supervoxel and for each of three behaviors: forward velocity, left rotational velocity, and right rotational velocity. The threshold for a significant correlation was set as p = 0.001 with a Bonferroni multiple comparison correction (2000 × 49 comparisons), giving a final p threshold of 1×10^−8^. Because supervoxels reduce the spatial resolution, we also calculated correlations of the individual voxels (x, y, z, t; 256, 128, 49, 30456). Indeed, we found that in this simple correlation analysis, the individual voxels produced a higher spatial resolution map. However, with so many comparisons, using a Bonferroni correction caused a very strong erosion of the map. Therefore, we used the approximate spatial coverage of the supervoxel map with its Bonferroni correction to set a reasonable p-value threshold for the individual voxel map (p = 1×10^−4^). This resulted in a map with the highest possible spatial resolution, while still using a principled means of setting the significance threshold.

#### Principal Component Analysis and Linear Modeling

Principal components were calculated using the superfly with supervoxels (supervoxel, z, t; 2000, 49, 30456). The matrix was reshaped as (feature, t; 98000, 30456), the covariance matrix was calculated, and an eigendecomposition was performed producing eigenvalues and eigenvectors. This was repeated for individual flies as well (not pooled into a superfly) to compare the differences (Figure 3G). Linear models were fit using principal components as features and a single behavioral variable as output. The number of features were sweeped to find the value that maximizes prediction accuracy (Figure 3F). Data was split into training and test sets and five-fold cross-validation was used to calculate the prediction accuracy on the held out test set (R^2^). A ridge penalty was employed to regularize the model and prevent overfitting.

#### Unique Variance Explained Modeling

Here, we used behavior to predict neural activity of each supervoxel independently (supervoxel, z, t; 2000, 49, 30456). For each supervoxel, we fit separate models with different combinations of input features: single behavior variables (forward velocity, left rotational velocity, right rotational velocity, and binary walking or not walking), all four behavior variables, or leave-one-out. We calculated the prediction accuracy for each model and for each supervoxel. To calculate the unique contribution of each behavior variable, we subtracted the prediction accuracy of the leave-one-out models from the all-four-variable model. All models were five-fold cross-validated and regularized with ridge regression.

#### Cross-Correlation Analysis

Given a matrix of timestamps for each slice of neural data acquisition across the superfly (z, vol_num; 49, 30456), windows of interpolation of behavior variables were created centered at each slice and volume number, with 20 ms steps extending 5 sec before and after. This resulted in a behavior matrix of (z, vol_num, interp_window; 49, 30456, 500). This matrix was then weighted by the neural activity for each supervoxel to produce a filter matching the interpolation window. This resulted in the equivalent of a cross-correlation filter for each supervoxel and for each behavior variable. We noticed these filters had a power peak in the frequency spectrum at the volume imaging rate (1.8 Hz), which we removed with a notch filter. Only the top 40% of responding filters were used for analysis, informed by the statistically significant fraction of the brain determined to correlate with behavior (previous correlation analysis).

#### GCaMP6f Deconvolution

The temporal dynamics were deconvolved from both the cross-correlation filters and the whole-brain spatiotemporal filters as follows. An experimental measurement of the GCaMP6f impulse response was used (100 ms to peak and 250 ms wide at half-max), taken from *Drosophila* early visual system neurons Tm3 and Mi1 (Yang et al., 2016). The measured kinetics were modeled by the formula ΔF/F = (1-e-t/4)*(-e-t/8) (Lütcke et al., 2013). This was expanded into a Toeplitz matrix and the deconvolved filters are given by the least-squares solution to y = Xb, where y is the measured convolved filter, X is the Toeplitz matrix, and b is the unknown deconvolved filter (Hansen, 2002).

## Supplemental Videos and Tables

**Supplemental Video 1. Brain-wide correlation with velocity space, coronal view, related to Figure 2**. Mean anatomical *Drosophila* brain colored by superfly correlations to velocity (n = 9 flies). Blue is correlation with forward velocity, green is correlation with left angular velocity, and red is correlation with right angular velocity. White lines indicate neuropil boundaries. Video moves from anterior to posterior. Top to bottom of any given frame is dorsal to ventral.

**Supplemental Video 2. Brain-wide correlation with velocity space, horizontal view, related to Figure 2**. Mean anatomical *Drosophila* brain colored by superfly correlations to velocity (n = 9 flies). Blue is correlation with forward velocity, green is correlation with left angular velocity, and red is correlation with right angular velocity. White lines indicate neuropil boundaries. Video moves from dorsal to ventral. Top to bottom of any given frame is anterior to posterior.

**Supplemental Video 3. Brain-wide unique contribution from velocity components, related to Figure 5**. Mean anatomical *Drosophila* brain colored by superfly unique variance explained of velocity components (n = 9 flies). Blue is unique contribution from forward velocity, green is unique contribution from left angular velocity, and red is unique contribution from right angular velocity. White lines indicate neuropil boundaries. Video moves from anterior to posterior. Top to bottom of any given frame is dorsal to ventral.

**Supplemental Video 4. Brain-wide temporal relationship between neural activity and velocity components, related to Figure 6**. Mean anatomical *Drosophila* brain colored by temporal relationship to each velocity component (n = 9 flies) (see Figure 6 for details). Neural activity in red voxels preceed velocity, white is contemporaneous with velocity, and blue lags behind velocity. Top brain relates to forward velocity, middle relates to left angular velocity, and bottom relates to right angular velocity. White lines indicate neuropil boundaries. Video moves from anterior to posterior. Top to bottom of any given frame is dorsal to ventral.

**Supplemental Table 1. Anatomical region abbreviations, related to Figures 2, 5, and 6**.

